# Chromatin dynamics identifies 78 genes at loci associated with elevated intraocular pressure and primary open-angle glaucoma

**DOI:** 10.64898/2026.03.12.711121

**Authors:** Nivedita Singh, Zachary Batz, Jayshree Advani, Milton A. English, Rupalatha Maddala, Ponugoti Vasantha Rao, Anand Swaroop

**Affiliations:** Neurobiology Neurodegeneration & Repair Laboratory, National Eye Institute, National Institutes of Health, Bethesda, Maryland, USA; Department of Ophthalmology, Duke University School of Medicine, Durham, NC 27710, USA

**Keywords:** Gene regulation, Non-coding genome, Capture Hi-C, 3D genome topology, Complex trait, Vision impairment

## Abstract

Primary open-angle glaucoma (POAG) is a chronic neurodegenerative disorder, and elevated intraocular pressure (IOP) represents the major, and only modifiable, risk factor for the disease. We modeled increased IOP by treating three primary human trabecular meshwork (TM) cell strains with dexamethasone, then generated a high-resolution map of promoter-centered chromatin contacts and regulatory modules to decipher how the genomic architecture and epigenetic state of disease-associated loci contribute to pathogenesis. We identify dynamic changes in chromatin compartments and looping, cis-regulatory elements and transcription factor hubs corresponding to altered transcriptional profile. By integrating GWAS-associated variants with dexamethasone-induced 3D chromatin landscape, we discovered 26 IOP- and 52 POAG- candidate causal genes, which belong to vesicle transport, TLR, MAPK and hippo-YAP signaling pathways. We also uncovered transcriptional regulatory role of 103 non-coding lead variants. Our studies provide a mechanistic framework of genetic complexity associated with ocular hypertension and POAG pathogenesis in addition to targets for therapies.

## INTRODUCTION

Glaucoma is a heterogeneous group of chronic multi-factorial optic neuropathies that lead to irreversible blindness, affecting nearly 80 million individuals worldwide (Tham et al., 2014). The disease manifests a complex clinical etiology, eventually resulting in loss of retinal ganglion cells and damage to the optic nerve. Primary open-angle glaucoma (POAG) is the most prevalent subtype that accounts for over 80% of glaucoma cases in United States (Allison et al., 2020; Friedman et al., 2004) and is characterized by increased resistance to aqueous humor (AH) outflow without anatomical changes to the iridio-trabecular meshwork angle (Weinreb et al., 2016). The key risk factors for POAG pathology include genetic susceptibility, advanced age, large vertical cup-to-disc ratio, and elevated intraocular pressure (IOP); the latter being the only known modifiable element (Craig et al., 2020; Shan et al., 2024; Weinreb *et al*., 2016). Current treatment options for POAG are limited, with primary focus on slowing disease progression by reducing IOP via surgical or pharmacological intervention (Conlon et al., 2017; Garg and Gazzard, 2020).

Both POAG and IOP are highly heritable (Wang et al., 2017), with at least 40% of POAG patients having a family history (Gramer et al., 2014; van Koolwijk et al., 2012). Genome-wide association studies (GWAS) across multiple ancestries have uncovered 226 IOP (Khawaja et al., 2018; MacGregor et al., 2018) and 359 POAG (Gharahkhani et al., 2021; Han et al., 2023) associated risk loci, with a significant overlap between the two (Xu et al., 2021). For example, IOP and POAG risk variants are co-localized near genes involved in regulation of cell adhesion (*ANGPT1*, *FERMT2*, *IGF1*), reactive oxygen species mitigation (*TXNRD2*, *KLF5*), actin skeleton organization (*AFAP1*, *GAS7*, *SPTNB1*, *ANTXR1*), and lipid metabolism (*ABCA1*, *PLCE1*, *DGKG*). However, nearly all lead GWAS variants as well as those in linkage disequilibrium (LD) are localized in the non-coding genome making it difficult to ascertain their functional significance (Schipper and Posthuma, 2022). Variants in cis-regulatory elements (CRE) can potentially modify the expression of distal genes hundreds of kilobases away. Regulatory effects are further modulated by tissue- and disease-specific epigenetic states and 3D chromatin arrangements; therefore, understanding the links between genetic variants and pathogenesis requires thorough investigation in the relevant biological context (Hamel et al., 2024).

IOP is determined by the balance between AH production and its outflow. Trabecular meshwork (TM), a mesenchyme-derived smooth muscle-like tissue, primarily regulates AH outflow resistance and is critical for maintaining IOP (Tamm, 2009). Prolonged use of glucocorticoids, a common treatment for ocular diseases, can elevate IOP by inducing widespread transcriptional changes in genes that are preferentially expressed in TM, leading to reduced phagocytosis, increased actin reorganization and formation of cross-linked actin networks (Bermudez et al., 2017; Kathirvel et al., 2022; Patel et al., 2023). Therefore, dexamethasone-treated TM cells have been widely used as a model for investigating increased IOP and glaucoma-like pathology (Medina-Ortiz et al., 2013; Raghunathan et al., 2015).

Here, we leverage this model system to decipher the gene regulatory networks associated with IOP and POAG and to understand the influence of GWAS-identified non-coding variants. We create a comprehensive promoter-centered genome topology map, with epigenetic landscape and transcriptional profiles, of three independent primary human TM cell strains before and after treatment with dexamethasone. Our analyses of multi-omic datasets identify altered 3D genome topology and epigenetic landscape of TM-expressed genes as well as transcription factor (TF) hubs associated with IOP-mimicking conditions. Integrations of chromatin looping and CREs with IOP and POAG GWAS variants have uncovered candidate causal genes and signaling pathways that contribute to IOP variations and POAG pathology. Our studies thus provide a genetic basis, and targets for therapies, of these debilitating chronic conditions.

## RESULTS

### Genome regulatory architecture of primary human TM cells

We first captured the transcriptome (total RNA-seq) alongside genome-wide identification of histone marks for transcriptionally active (H3K27ac, H3K4me1/2/3) and repressed regions H3K27me3/H3K9me3), chromatin accessibility(Buenrostro et al., 2015), and CTCF binding of three independent primary human TM cell strains from healthy donor eyes expressing juxtacanalicular region marker genes (Fig. 1A, Supplementary Fig. 1A). As predicted, genes with an enrichment of activating histone modifications exhibit higher expression, whereas repressing marks are localized primarily to genes showing low expression (Supplementary Fig. 1B). To annotate TM cell-preferred CREs, we organized the chromatin accessibility, CTCF binding, as well as active/repressive histone marks into 13 epigenetic states using a Hidden Markov model (Fig. 1B) (Ernst and Kellis, 2017). We note that the regions classified as active promoters are concentrated in the 5’-UTR and/or near transcription start site (TSS) of genes. Enhancers are more diffusely recognized across the genome, whereas markers for active transcription tend to be elevated across gene bodies and especially near the 3’-region of genes (Fig. 1B).

**Fig. 1.**
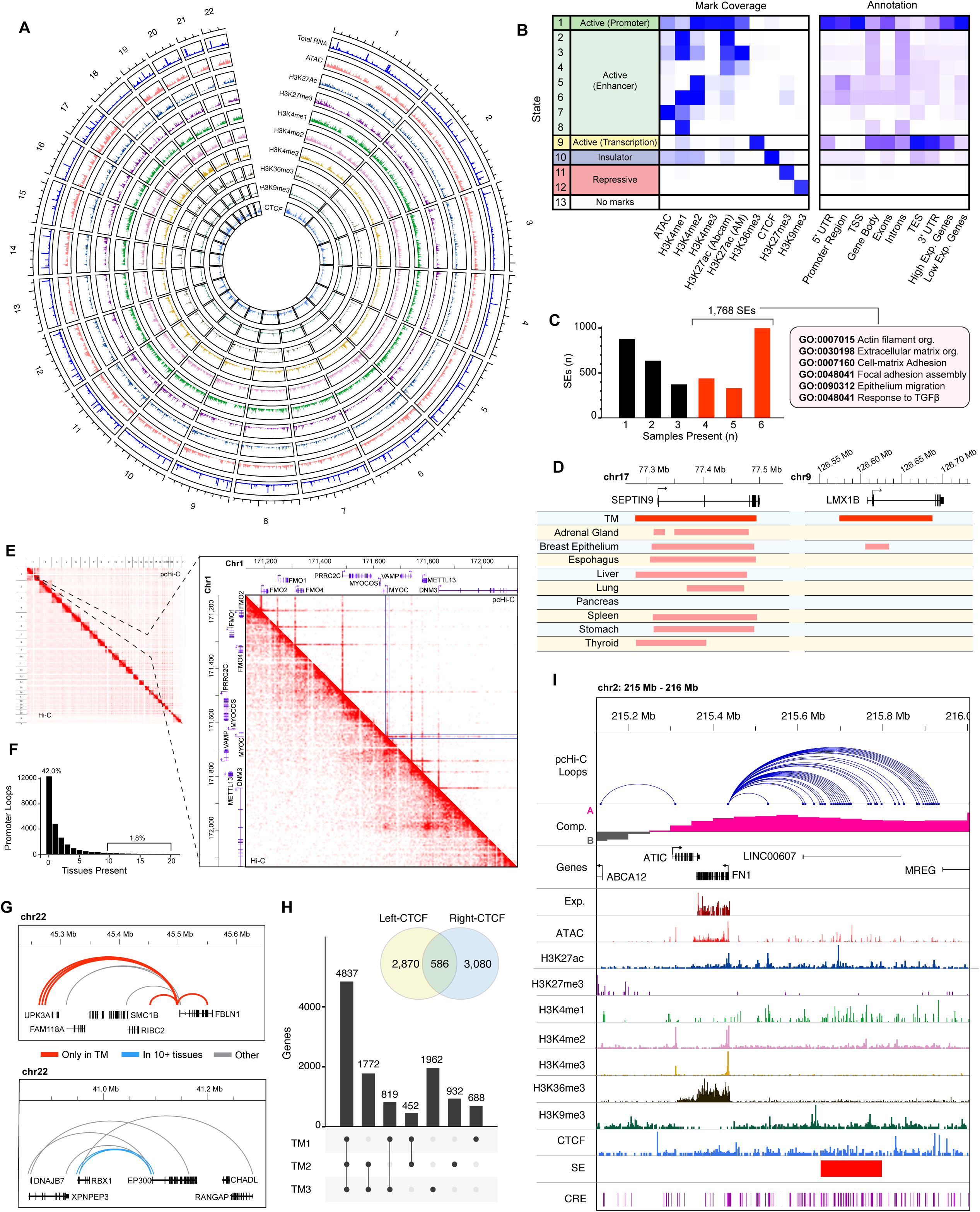
Summary of multi-omic resource derived from primary trabecular meshwork cells. **[A]** Overview of total transcription, epigenetic marks, and genome accessibility across trabecular meshwork (TM) autosomes. **[B]** Relative density of epigenetic marks called across chromatin states (left) and annotation associated with these regions (right). **[C]** Count of shared super enhancers (SEs) identified using six different sets of H3K27ac peaks (2 H3K27ac antibodies used on each sample). The 1,768 SEs that appear in at least half of these sets (red bars) were retained; select GO terms enriched among genes overlapping these SEs are shown (full GO term enrichment in Supplementary Fig. 1C). **[D]** Example of SEs observed in nearly all tissues (SEPTIN9 locus, left) and another rarely found outside TM (*LMX1B,* right). **[E]** Traditional (lower triangle) and promoter-capture (upper triangle) Hi-C contact maps across entire genome (left); Inset region highlights loop enrichment in promoter-capture Hi-C data near the *MYOC* gene. **[F]** Proportion of TM promoter-interacting loops observed in promoter-capture HiC from 20 tissues(Jung *et al*., 2019). **[G]** Example of TM-specific promoter loops at the *FBLN1* locus (upper) and universally observed promoter loops at the *EP300/RBX1* locus (lower). **[H]** Count of loops with CTCF peaks at one or both anchors (venn diagram inset); total genes in contact with CTCF-anchored loops in each combination of samples. **[I]** Epigenetic marks, expression, accessibility, and genome topology at the *FN1* locus.

Our analysis uncovers 1,768 super enhancers (SEs), which overlap genes enriched for canonical TM cell functions such as extracellular matrix organization, cell adhesion, TGFβ response, and actin filament assembly (Fig. 1C, Supplementary Fig. 1C). These regulatory regions are also associated with high gene expression (Fig. 1C, Supplementary Fig. 1D). A comparison of TM SEs, identified here, with 1.17 million SEs cataloged across 1,739 human samples(Wang et al., 2023) reveals that the average TM SE is only present in less than 15% of non-TM samples (Supplementary Fig. 1E). Few SEs appear to be nearly universal; e.g., the SE overlapping the cell cycle regulatory gene *SEPTIN9* is present in 1,603 samples (92.1%) including TM (Fig. 1D). However, nearly a third of SEs (31.5%) are detectable in fewer than 5% of other tissues; e.g., the TM SE over *LMX1B*, a gene linked to actin cytoskeleton maintenance (Burghardt et al., 2013), is observed in 36 samples (2.1%) reflecting specificity of its regulatory action (Fig. 1D).

We then performed promoter capture HiC (pc-HiC) (Schoenfelder et al., 2018) to identify promoter-centered chromatin contacts. Our pc-Hi-C analyses of three independent human TM cell strains identified 30,375 loops; of these, 26,495 (87.2%) loops overlap promoters including the key TM genes, such as *MYOC*, thereby revealing interactions among distal and proximal CREs (Fig. 1E). We then evaluated the specificity of these promoter-centered contacts by comparing promoter-interacting loops from TM cells to those observed across 20 human tissues (Jung et al., 2019). Notably, 42% of promoter loops identified in TM cells are not detected in all other tissues examined (Fig. 1F). As an example, *FBLN1*, a highly expressed gene encoding a major component of extracellular matrix (Li et al., 2023), has several promoter-centered loops unique to TM (Fig. 1G). In contrast, only 1.7% of TM promoter loops are observed in at least half of the tissues. We identify 534 genes that share multiple promoter loops across 10 or more tissues including TM, and these genes are enriched for constitutive functions such as cell cycle regulation, chromatin organization, and DNA repair (Supplementary Fig. 1F). For example, promoter-promoter contact between *RBX1*, a gene regulating the protein turnover (Hershko and Ciechanover, 1998), and the histone acetyltransferase (Ogryzko et al., 1996) *EP300*, is detected in 15 of the 20 tissues examined (Fig. 1G).

Chromatin contacts are anchored by CTCF, an architectural protein that mediates long-range interactions to regulate gene expression (Dehingia et al., 2022). We show that 6,536 promoter loops are associated with at least one CTCF-bound foot and that 586 loops have CTCF peaks on both ends. In total, 4,837 genes possess CTCF-anchored promoter loops in all three TM samples (Fig. 1H). These genes were enriched for pathways such as growth, organelle localization, and actomyosin organization (Supplementary Fig. 1G).

Taken together, this multi-omic framework provides insights into regulation of genes associated with TM function. We specifically highlight the chromatin context of fibronectin-1 (*FN1*; Fig. 1I), a key component of the extracellular membrane of TM cells (Medina-Ortiz *et al*., 2013). The *FN1* gene is highly expressed in TM and localized to compartment A. Its promoter is enriched in activating histone marks (H3K27ac and H3K4me3) and deficient in repressive marks (H3K27me3, and H3K9me3). The active transcription of *FN1* is consistent with a broad peak of H3K36me3. We also uncover 35 promoter-interacting loops, many of which contact a SE located ∼200kb upstream of TSS.

### Dexamethasone reshapes the transcriptional and epigenetic landscape of TM cells

Treatment of TM cells with dexamethasone, a glucocorticoid commonly used to mimic IOP elevation *in vitro* (Overby and Clark, 2015; Watanabe et al., 2021; Yemanyi et al., 2020), results in global changes in transcriptome; Principle component analysis shows that dexamethasone treatment accounts for ∼13% of the total variation in expression (Fig. 2A, Supplementary Fig. 2A and 2B) with 3,423 differentially expressed genes (1,527 showing higher, 1,894 lower expression; Fig. 2B). Given that dexamethasone’s effect is mediated via glucocorticoid receptors (GR), over half (51.4%) of the 108 putative glucocorticoid target genes (Choi et al., 2023) are differentially expressed following the treatment (Supplementary Fig. 2C).

**Fig. 2.**
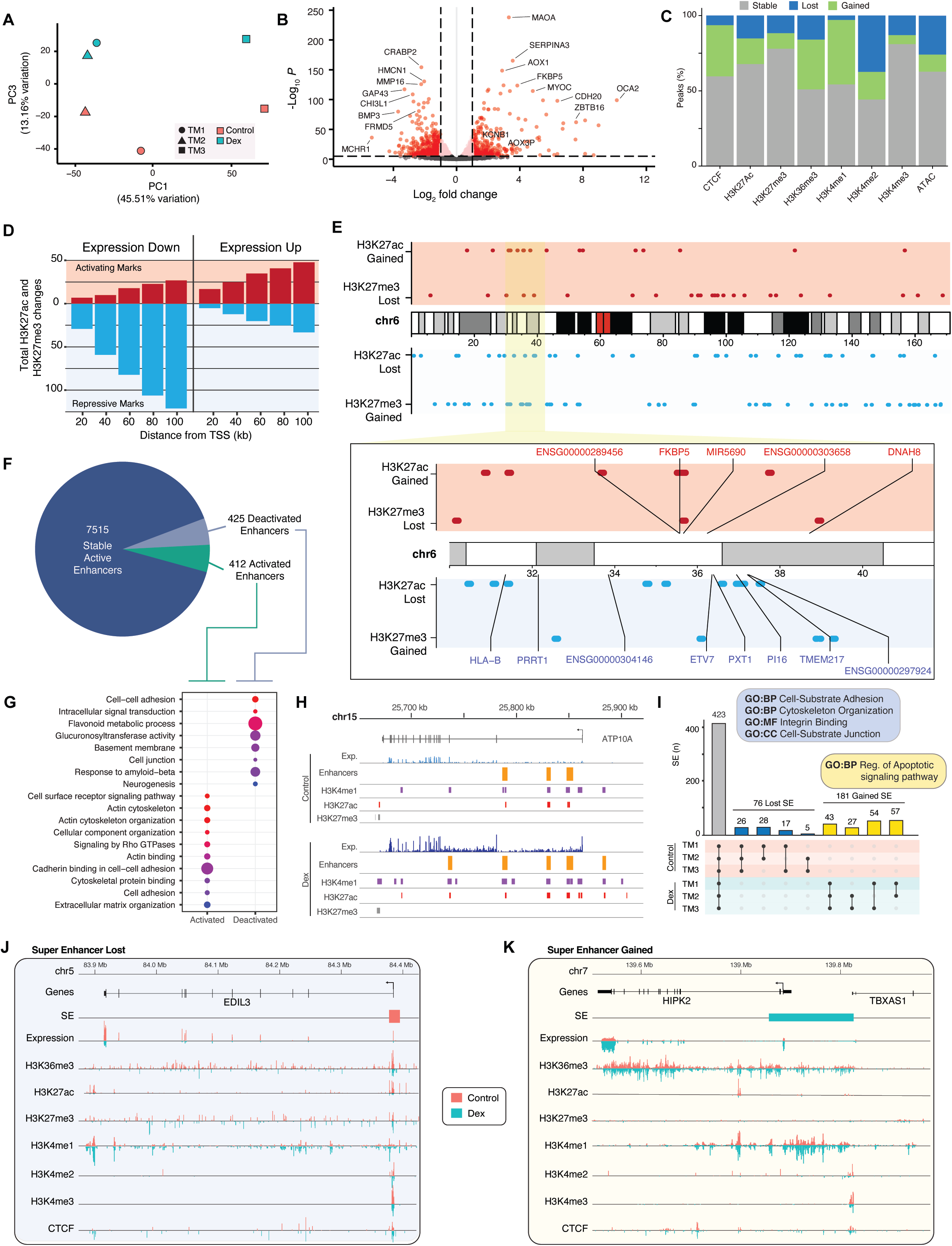
Dexamethasone broadly alters transcriptional and epigenetic state of trabecular meshwork cells. **[A]** Summary of global transcriptional state for trabecular meshwork (TM) samples by control line and treatment status for PC1 and PC3. Dexamethasone-induced changes in **[B]** the expression of individual genes and **[C]** peaks for CTCF, ATAC-seq, and histone marks. **[D]** Count of activating epigenetic changes (i.e., gain of H3K27ac; loss of H3K27me3) and repressive epigenetic changes (i.e., gain of H3K27me3; loss of H3K27ac) by proximity to transcription start sites (TSS) for differentially expressed (DE) genes after dexamethasone treatment. **[E]** Map of H3K27ac and H3K27me3 peak changes across chromosome 6. Activating epigenetic changes are marked in the red panels; repressive epigenetic changes are marked in the blue panels. Inset zoom highlights peak changes in the region surrounding the *FKBP5* locus. Gene names in red have >1 log_2_fold change after dexamethasone treatment; gene names in blue have <-1 log_2_fold change after dexamethasone treatment. **[F]** Summary of changes in shared active enhancers after dexamethasone treatment. Enhancer state identification was performed as described in Methods. **[G]** Select GO terms enriched among genes overlapping enhancers which either are activated or deactivated following treatment. **[H]** Enhancers, expression, and histone marks at the *ATP10A* locus with and without dexamethasone treatment. **[I]** Super enhancer (SE) changes by TM line and treatment. Inset blue and yellow boxes contain select GO terms enriched among DE genes overlapping lost and gained SEs, respectively. Example loci with **[J]** lost (*EDIL3*) and **[K]** gained (*HIPK2*) SEs.

Further analysis of the total RNA-seq data predicts 4,152 expressed enhancer RNAs (eRNAs; Supplementary Fig. 2D); 51 of these are differentially expressed after dexamethasone treatment (Supplementary Fig. 2E). Curiously, one the predicted eRNA and its proximal gene *ALOX15B* both show significantly augmented expression after dexamethasone (Supplementary Fig. 2F). Increased expression of *ALOX15B* has been noted during early stages of ocular hypertension in mouse models (Wils et al., 1999).

Histone modifications, chromatin accessibility, and CTCF binding profiles demonstrate widespread epigenetic alterations following dexamethasone treatment (Fig. 2C). Overall, chromatin accessibility is reduced with loss of 25.8% of peaks, including 2,628 overlapping gene promoters and SEs (Supplementary Fig. 2G). Furthermore, we observe a significant reduction of H3K4me2 marks that are associated with tissue-specific TF binding at enhancers and edges of promoters (Shah et al., 2018; Wang et al., 2014). In contrast, dexamethasone-treated cells exhibit elevated CTCF, H3K36me3, and H3K4me1 binding that are linked to chromatin insulation, active transcription over gene body, and promoters, respectively.

Relatively few changes are observed in the absolute number of H3K27ac and H3K27me3 peaks at active enhancers and repressors, respectively. Yet, the changes in distribution of these marks coincide with altered gene expression patterns. Genes showing lower expression after dexamethasone treatment tend to reside near regions shifting towards a repressive epigenetic state (i.e., gain of H3K27me3 and/or loss of H3K27ac), whereas those presenting higher expression are colocalized with activating epigenetic changes (i.e., gain of H3K27ac and/or loss of H3K27me3; Fig. 2D). As an example, the genomic region surrounding the *FKBP5* gene, a functional regulator of the glucocorticoid receptor complex (Binder, 2009), is enriched for activating epigenetic marks, concurrently gaining H3K27ac and losing H3K27me3 (Fig. 2E). *FKBP5* and other genes at this locus demonstrate significantly increased expression following dexamethasone treatment. In contrast, significant reduction in expression of five genes at the genomic region ∼1Mb away is associated with the loss of multiple H3K27ac peaks (Fig. 2E). Large scale chromatin reorganization at the *FKBP5* locus in response to glucocorticoid treatment has also been reported in mouse and human macrophages (Jubb et al., 2017).

We further annotated enhancers based on the activation state; 8,342 enhancers are present in both treatment groups and active in at least one (Fig. 2F). Approximately 10% of these enhancers switch between an active and non-active (i.e., poised, primed, or inactive) state after treatment. Dynamic enhancers overlap genes that are enriched for ECM organization, actin morphology, cell adhesion and Rho GTPase signaling (Fig. 2G); all known to regulate IOP homeostasis in TM cells (Pattabiraman et al., 2013; Soundararajan et al., 2021; Vranka et al., 2015). We specifically note the gain of H3K27ac peaks and switch to an active state for two primed enhancers at a POAG locus near *ATP10A*, which exhibits increased expression after dexamethasone treatment of TM cells (Fan et al., 2008).

Over one-third of TM SEs (37.8%) are gained or lost following dexamethasone treatment (Fig. 2I). Gained SEs are enriched for apoptotic genes, whereas lost SEs are for genes associated with cytoskeletal and cell binding functions. For example, one SE overlapping the promoter/TSS region of *EDIL3* was lost with a concurrent decrease in gene expression (−0.61 log_2_ fold change). At this locus, we detect reduced H3K27ac and H3K36me3 as well as enhanced H3K4me1 and H3K27me3 marks, consistent with a shift from an active to a poised enhancer. *EDIL3* encodes an integrin binding protein that is abundant in the TM extracellular vesicles and reduced in glaucomatous TM cells (McDonnell et al., 2023). In contrast, the gain of SE at the promoter/TSS region of *HIPK2* is accompanied by its augmented expression (+1.7 log_2_ fold change) after dexamethasone treatment. *HIPK2* is a serine/threonine kinase that is a positive regulator of fibrosis (Garufi et al., 2023), which can potentially reduce aqueous humor outflow and cause elevated IOP.

### Dexamethasone alters 3D genome topology in TM cells

Dexamethasone-induced transcriptome changes can be mediated, at least in part, by remodeling of chromatin compartments and promoter-interacting loops. We called the compartments at 50kb resolution and identified significant shifts in 493 compartment windows, with displacement towards a heterochromatic B compartment state observed more frequently (Fig. 3A). Though only 0.7% of the total, these dynamic regions appear to be hotspots for transcriptomic and epigenetic changes (Fig. 3B). Windows moving towards euchromatic A compartment reveal high gene expression, along with increased H3K27ac, H3K4me1 and H3K36me3 signals, as well as a decrease in H3K27me3 marks. In contrast, shifts towards B compartment correspond to lower gene expression and reduced marks for H3K27ac, H3K4me1 and H3K4me3. Both classes of compartments exhibit lower levels of H3K4me2. We note a similar reduction in H3K4me2 even in genomic regions showing no change in chromatin compartments (Supplementary Fig. 3A), suggesting that overall signal for this mark is lower after dexamethasone treatment.

**Fig. 3.**
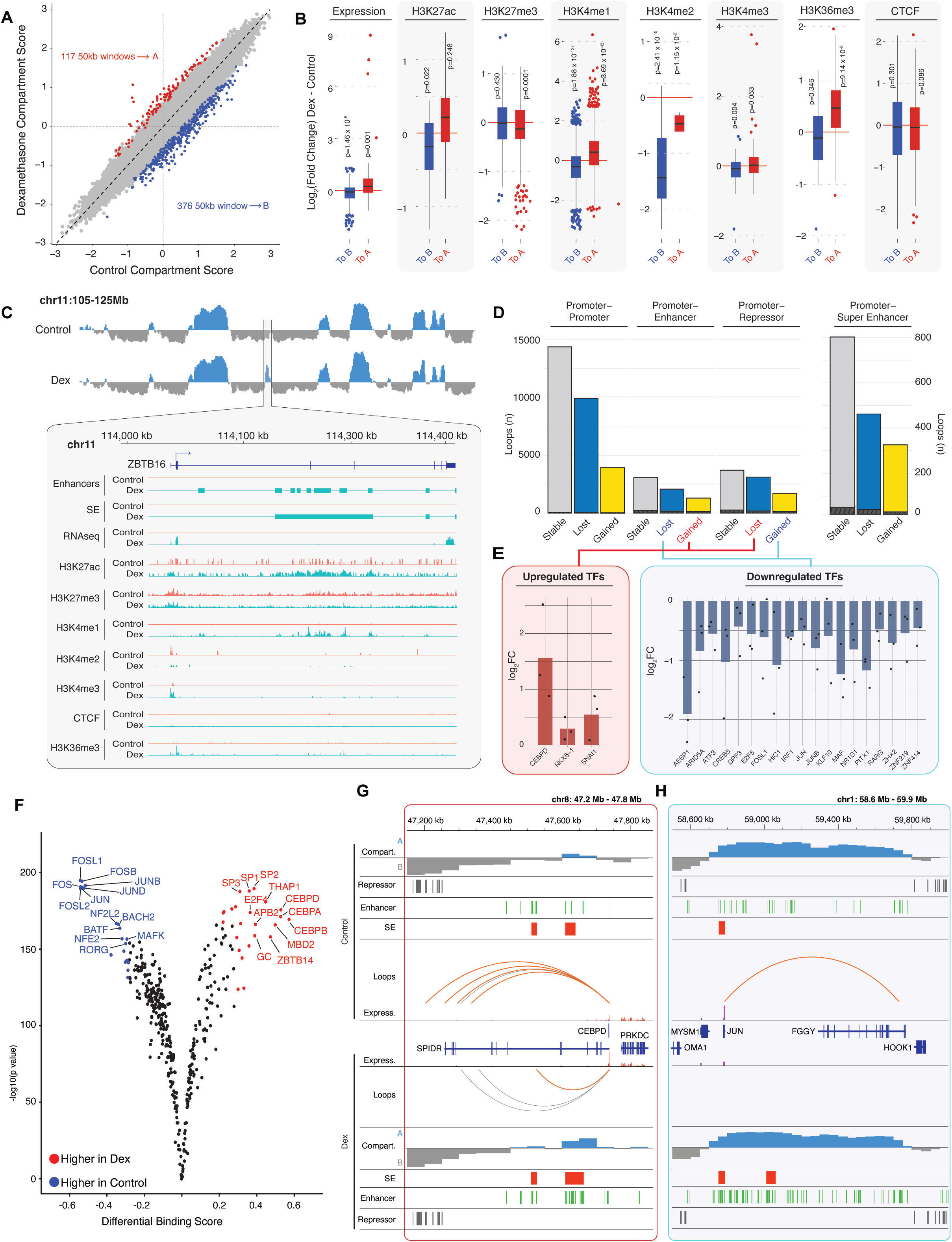
Dexamethasone treatment modifies higher order chromatin structure in TM cells. **[A]** Compartment score changes for each 50kb genomic window, windows with significant shifts towards A or B compartment are highlighted in red and blue, respectively. **[B]** Changes in transcription, histone marks, and CTCF signal within changed compartments. Box plots show interquartile range (IQR) and whiskers represent 1.5x the IQR. One-way t-tests were used to determine each distribution differs significantly from zero (red line). **[C]** A and B compartments identified on chromosome 11 (blue and gray for A and B compartments, respectively). Inset zoom highlights compartment change from B to A compartment at the *ZBTB16* locus following dexamethasone treatment. Red and blue lines show signal from control and dexamethasone-treated TM cells, respectively. **[D]** Count of promoter-interacting loops present in control samples only (lost), dexamethasone treated samples only (gained), or in both (stable). Hashes indicate transcription factor (TF) promoters within each bar. **[E]** Differentially expressed TFs identified with activating changes (i.e., gained promoter-enhancer contact and/or lost promoter-repressor contact) or repressive changes (i.e., lost promoter-enhancer contact and/or gained promoter-repressor contact). Dots represent expression changes in individual TM strains, bars represent the mean of expression change observed across all samples. **[F]** Predicted shifts in TF binding based on ATAC footprinting analysis (see Methods). Red and blue dots indicate TFs with significantly more or less bound regions, respectively, after dexamethasone treatment. Promoter-interacting loops, compartments, expression and epigenetic state of **[G]** *CEBPD* and **[H]** *JUN* loci. Red loops are gained, and yellow loops are lost after dexamethasone treatment.

One of the dynamic regions includes ZBTB16, a Zn-finger TF that interacts with histone deacetylase and is reported to show dramatically higher expression in TM cells in response to dexamethasone (Rozsa et al., 2006). A shift from B to A compartment and in chromatin landscape at the *ZBTB16* gene region coincides with augmented transcription from <1 CPM in control samples to >150 CPM after dexamethasone treatment (Fig. 3C). We observe increased signals for H3K27ac, H3K4me1, H3K36me3, and H3K4me3 as well as emergence of multiple enhancers and a SE overlapping the gene body. Furthermore, we detect the loss of looping upstream of *ZBTB16* likely due to the appearance of a TAD boundary (Supplementary Fig. 3B). Notably, dexamethasone treatment of kidney podocytes similarly leads to the emergence of a SE that overlaps *ZBTB16* driving increased acting filament synthesis and improving cytoskeletal stabilization (Wang et al., 2021).

In TM cells, dexamethasone treatment alters 23,397 promoter-interacting loops, including 1,350 loops that relate to SEs and 557 with promoters of TFs (Fig. 3D), which can serve as hubs for IOP and POAG gene networks. Three TFs corresponding to activating promoter loop changes (i.e., gain of loops to enhancers or loss of loops to repressors) exhibit higher expression, whereas 19 TFs showing lower expression show gain of loops to repressors and/or loss of loops to enhancers (Fig. 3E). Next, we assessed whether changes in the expression of TFs correspond to changes in TF binding using ATAC footprinting (Bentsen et al., 2020). Many TFs indeed reveal dramatically altered binding scores after dexamethasone (Fig. 3F); these include CEBP, JUN and FOS family proteins (Fig. 3G, 3H). The expression of *CEBPD* increases concurrently with the gain of contact between the promoter region and an intronic SE, loss of contact with intergenic repressors, and a shift towards A compartment (Fig. 3G). In contrast, AP-1 constituent TFs have reduced contact with distal CREs and exhibit lower expression (Fig. 3E,3H), possibly due to the loss of an enhancer-rich region upstream of JUN following the dexamethasone treatment (Fig. 3H). Interestingly, 177 genes with CEBPD-bound promoters specific to dexamethasone are enriched for MAPK signaling as well as regulation of necrosis and extracellular matrix organization (Supplementary Fig. 3C and 3D). In contrast, JUN-bound promoters of 438 genes that are specific to control samples are enriched for regulation of WNT and RAP1 signaling, and interleukin 18, among others (Supplementary Fig. 3E and 3F).

### Dexamethasone causes shifts in TM 3D genome topology, associated with global downregulation of TNF signaling

We then carried out separate enrichment analyses for gene sets exhibiting changes in transcription, epigenetic marks, compartmentalization, or distal contact patterns (see Methods), and intersected these analyses to distinguish biologically meaningful units that are altered across multiple levels of regulation (Fig. 4A). Integrated analyses of multi-omic datasets reveal widespread and concordant epigenetic and transcriptional changes in genes associated with key TM functions including focal adhesion, actin cytoskeleton, ECM structure and fibronectin binding (Fig. 4A). TNF and AGE-RAGE signaling are the two pathways most consistently identified across these analyses (Fig. 4B, S4A, S4B).

**Fig. 4:**
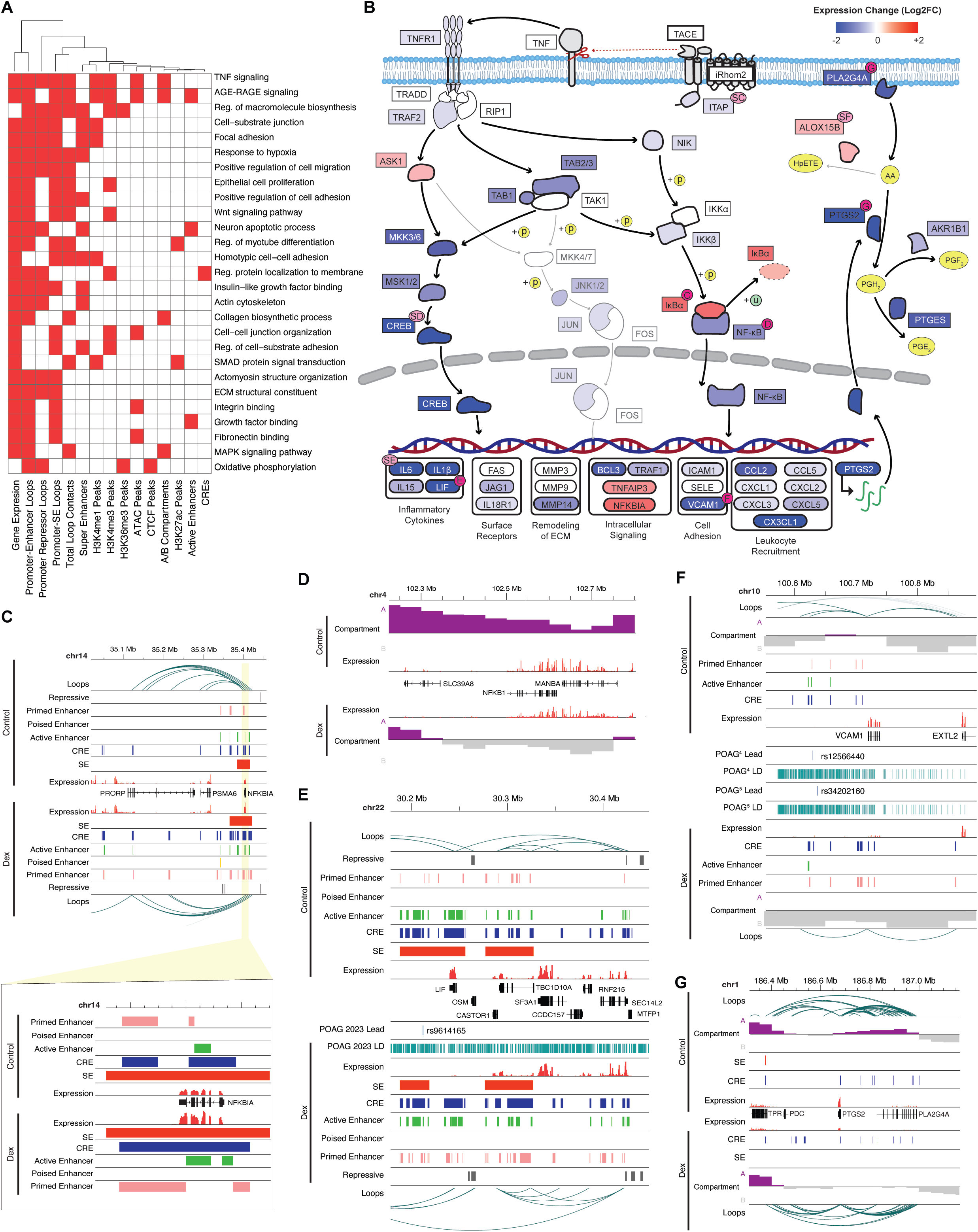
TNF signaling pathway is broadly repressed after dexamethasone treatment. **[A]** Overview of enriched terms in dexamethasone-treated cells. Columns represent gene sets altered by dexamethasone treatment (e.g., genes that changed expression, genes gaining or losing promoter-enhancer loops, etc); rows represent individual GO or KEGG terms. Red squares indicate a given term is enriched within a gene set. All terms enriched in at least 4 gene sets are shown. **[B]** Expression changes in the TNF signaling pathway. Pink circles indicate genes where epigenetic/chromatin changes and GWAS variants are further highlighted in additional panels including **[C]** *NFKBIA,* **[D]** *NFKB1*, [**E]** *LIF,* **[F]** *VCAM1*, and **[G]** *PTSG2*-*PLA2G4A*. Additional genes are shown in **Supplemental Fig. 4**.

Coordinated downregulation of TNF signaling is evident with corresponding changes in epigenome marks of multiple TFs and their gene targets (Fig. 4B). Expression of the gene encoding iTAP (*FRMD8*), which is required to release TNF from the cell membrane and initiate TNF signaling (Oikonomidi et al., 2018), is reduced by 25% (FDR = 0.004), gaining a repressive mark directly over the promoter region after dexamethasone (Supplementary Fig. 4C). We also note that promoter-interacting loops observed exclusively in the dexamethasone-treated samples contact the genomic region including rs1346, a lead SNP identified in POAG-GWAS (Supplementary Fig. 4C) (Gharahkhani *et al*., 2021; Han *et al*., 2023).

Dexamethasone response results in ∼30% lower expression of the *NFKB1* gene encoding NFkb and 2-fold higher expression of the *NFKBIA* gene that encodes its inhibitor. The regulatory mechanisms orchestrating these changes appear diverse. Enhanced expression of *NFKBIA* may be driven by distal enhancer CREs gained in regions of loop contact to the promoter as well as by the activation of an enhancer directly over its TSS (Fig. 4C). In contrast, repression of *NFKB1* coincides with a local shift to heterochromatic B compartment after dexamethasone (Fig. 4D). Expression of *CREB5*, encoding another TF in this pathway, is reduced by almost 50%, likely due to the loss of loop contact with an upstream enhancer region (Supplementary Fig. 4D).

Downstream regulatory targets of these repressed TFs include inflammatory cytokines such as *LIF* and *IL6* (Banner et al., 1998; Uciechowski and Dempke, 2020). *LIF* expression is reduced over 80% and correlates with some combination of deactivation of enhancers directly over the TSS region, loss of contact between the promoter and an upstream cluster of active enhancers, and the loss of SE that directly overlaps the gene in control conditions (Fig. 4E). We further note that the lost SE contains rs9614165, a lead SNP for POAG (Han *et al*., 2023). Changes in the *IL6* gene are similar; its expression is reduced by 88% and a SE in contact with the promoter region via looping is lost in response to dexamethasone (Supplementary Fig. 4E). Additional regulatory targets include *VCAM1*; this gene encodes a cell adhesion molecule and its expression is decreased ∼70% (Fig. 4F) as the locus shifts towards B compartment, loses distal enhancers, and exhibits reduced looping to an upstream CRE near POAG lead SNPs(Fig. 4F) (Gharahkhani *et al*., 2021; Han *et al*., 2023).

We also identify changes in the TNF target gene, *PTGS2* (also called *COX2*) (Fig. 4G). *PLA2G4A* and the adjacent *PTGS2* both encode enzymes necessary for prostaglandin synthesis (Clark et al., 1991; Vane et al., 1998) and their expression is significantly reduced following dexamethasone treatment. The locus containing these genes shifts from A to B compartment, and *PLA2G4A* loses a CRE near its promoter post-dexamethasone. The *PTGS2* gene encoding an enzyme that catalyzes arachidonic acid to prostaglandin H_2_ (Vane *et al*., 1998) loses a distal SE connected to the promoter region by looping. This reaction is negatively correlated with the conversion of arachidonic acid into hydroperoxy eicosatetraenoic acids by ALOX15B (Meng et al., 2018). The TSS region of *ALOX15B* loses a repressive mark and gains an enhancer while augmenting expression from <0.1 CPM in control to >2.5CPM after dexamethasone treatment. Prostaglandin H_2_ is further converted on to PGF_2a_ catalyzed in part by the product of AKR1B1 (Bresson et al., 2011) and the expression of this gene is reduced by ∼25% (Fig. 4B). Analogs of PGF_2a_ are used frequently to treat elevated IOP underscoring the importance of changes in this biosynthetic pathway for glaucoma pathogenesis.

### 3D genomic architecture links IOP- and POAG- GWAS associated non-coding variants to cognate gene promoters

We then sought to connect promoter loops and SEs with lead GWAS variants identified for IOP (Khawaja *et al*., 2018; MacGregor *et al*., 2018) and POAG (Gharahkhani *et al*., 2021; Han *et al*., 2023) as well as variants in LD (Genomes Project et al., 2015). For each study, we identify loops connecting gene promoters to either lead GWAS SNPs or variants in LD in each of our three chromatin loop sets [i.e., resource (Fig. 1); control; and dexamethasone-treated (Figs 2-3)]. We identified a total of 26 and 52 lead variants in contact with gene promoters for IOP and POAG, respectively (Fig. 5A, Table S1). Target genes for these lead variants are enriched for pathways such as vesicle-mediated transport, Toll-like Receptor, MAPK and Hippo signaling. This topological data provides candidate genes for intergenic variants; for example, the promoter for *LRIG1* forms extensive loop contacts with a region more than 300kb away that contains SNPs identified in both IOP and POAG GWAS datasets (Fig. 5B). Next, we assess the tissue-specificity of loops connecting lead SNPs to gene promoters in comparison to publicly available promoter capture Hi-C data from 20 tissues(Jung *et al*., 2019). Only one SNP-promoter loop was observed in a majority of tissues (rs1782976-*APP*), whereas more than half were unique to TM cells (IOP: 55.2%; POAG: 52.6%; Fig. 5C, Supplementary Fig. 5A). For example, the SNP rs17752199 is proximal to *PKHD1* but forms long range contacts with the promoter of *TFAP2B* (Fig. 5C). In mice, a deletion from intron 2 to the 3’UTR of *Pkhd1* reduces *Tfap-2b* expression and results in elevated IOP and RGC death (Ishimoto et al., 2026). Our results are consistent with the hypothesis that disruption of local chromosomal architecture interferes with *TFAP2B* expression in human TM.

**Fig. 5:**
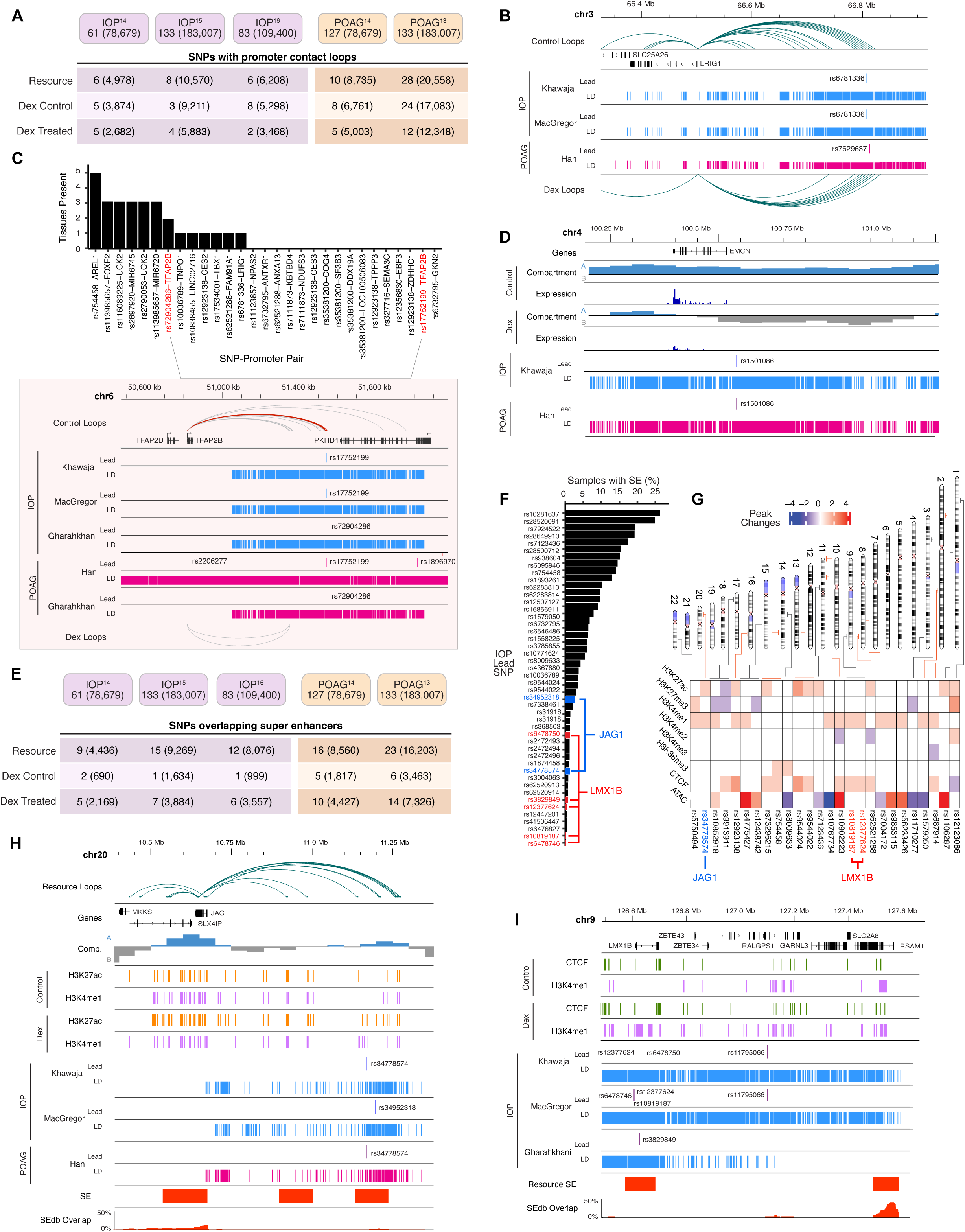
IOP and POAG GWAS variants are connected to target genes via promoter loops and super-enhancers (SEs). **[A]** Count of IOP and POAG different GWAS lead variants and variants in linkage disequilibrium (LD variants; shown in parenthesis) contacting genes via promoter loops in TM cells **[B]** Loops in control and dexamethasone treated samples connect IOP and POAG lead variants to the *LRIG1* promoter. **[C]** Count of tissues(Jung *et al*., 2019) overlapping each IOP lead SNP-Promoter pair observed in TM cells. Inset panel shows loop connecting rs17752199 and *TFAP2B* in red. **[D]** Lead GWAS SNPs and compartment changes after dexamethasone near *EMCN*. **[E]** Count of IOP and POAG lead variants and variants in LD (shown in parenthesis) overlapping a SE in each dataset. **[F]** Percent of samples from SEdb 2.0 (n=1,739) overlapping each super enhancer containing a lead IOP SNP. SNPs of interest near *JAG1* and *LMX1B* are indicated in blue and red, respectively. **[G]** Number of peaks gained or lost following dexamethasone treatment within 5kb of lead IOP SNP. Ideogram on the top indicates the position of each SNP. SNPs of interest near *JAG1* and *LMX1B* are indicated in blue and red, respectively. Super enhancers, loops, and changes in histone marks near **[H]** *JAG1* and **[I]** *LMX1B*.

We further ascertained 9 and 18 IOP and POAG lead SNPs, respectively, overlapping loci that change compartments after dexamethasone treatment; these include five SNPs identified in both IOP- and POAG-GWAS. For example, the region containing *EMCN* and the lead SNP rs1501086 switches from A to B compartment in response to dexamethasone treatment and the *EMCN* gene has reduced expression by approximately half (Fig. 5D). *EMCN* is a marker gene for endothelial cell types (Liu et al., 2001) and inhibits focal adhesion between cells and ECM (Kinoshita et al., 2001), pointing to a potential role in regulating AH outflow in TM tissue.

Next, we identified a maximum of 15 and 23 lead variants overlapping SEs from IOP and POAG GWAS sets, respectively (Fig. 5E). On average, SEs containing SNPs are relatively tissue-specific, with just 6.4% of external, non-TM samples(Han *et al*., 2023) overlapping SEs with IOP lead variants and 9.4% with POAG lead variants (Fig. 5F, S5B). For each SNP, we quantify changes in histone, ATAC, and CTCF peaks within 5kb regions (Fig. 5G, S5C). Lead SNPs near *JAG1* and *LMX1B* reside in TM-specific SEs and are co-localized with extensive epigenetic changes induced by dexamethasone treatment. *JAG1* maintains promoter loops to an A compartment region ∼500kb away containing a SE observed in just 0.8% of non-TM samples. The distal SE region contains lead SNPs identified in both IOP and POAG studies and gains H3K27ac and H3K4me1 peaks for both after the addition of dexamethasone (Fig. 5H). We similarly detect multiple lead IOP SNPs near *LMX1B* (Fig. 5I) and identify a SE overlapping the gene promoter, which is present in just 0.2% of non-TM samples. After dexamethasone, the SE locus shows higher H3K4me1 and CTCF binding consistent with a change in promoter regulation. *LMX1B* encodes a LIM-homeodomain protein required for TM formation (Liu and Johnson, 2010). In humans, *LMX1B* mutations cause nail-patella syndrome (Dreyer et al., 1998), a disease linked to elevated risk of glaucoma (Farley et al., 1999), whereas heterozygous *Lmx1b* mutations results in elevated IOP and glaucoma in mice (Cross et al., 2014).

## DISCUSSION

Elevated IOP and hundreds of non-coding genetic variants are robustly linked to POAG susceptibility. There is an unmet need to integrate POAG and IOP associated variants in the non-coding genome to transcriptional profile of the coding genome and identify underlying POAG genes, as reported in other complex traits (Nott et al., 2019; Schoenfelder and Fraser, 2019). Our study highlights the importance of elucidating cell-type specific transcription regulatory modules, identifying CREs and promoter-centered chromatin contacts unique to TM. In elevated IOP-mimicking conditions, we uncover highly orchestrated and dysregulated changes across genome architecture, epigenetic landscape, transcription factor hubs and transcriptional patterns including in established disease-associated genes. For example, we reveal regulatory mechanisms likely controlling TM-specific MYOC transcriptional circuits in homeostasis and POAG; mutations in this gene are associated with 2-4% of adult-onset POAG and 10% juvenile open angle glaucoma (Saccuzzo et al., 2023). Intriguingly, multiple genes required for arachidonic acid (AA) metabolism undergo coordinated changes including *ALOX15B*, which mediates the inflammation reducing effect of glucocorticoids by increasing the biosynthesis of specialized mediators (Rao et al., 2023). Yet, TM inflammation has been hypothesized to contribute to POAG pathogenicity (Baudouin et al., 2021; Cela et al., 2021) in apparent contrast to increased anti-inflammatory activity driven by elevated *ALOX15B* expression after dexamethasone. Recent results from rat glaucoma models have demonstrated that the IOP lowering effects of the glaucoma drug Latanaprost are both mediated by *ALOX15B* upregulation and PTGS2-driven prostaglandin synthesis pathway (Mathew et al., 2025). Our studies uncover widespread downregulation of this key pathway in elevated IOP-mimicking conditions and reveal regulatory hubs likely underpinning this pathogenic repression.

### Dysregulation of transcription factor targets

Footprinting analysis has allowed mapping of dynamic DNA-protein binding of CEBPD and JUN. Effector genes of CEBPD, predicted to have increased binding after dexamethasone, are enriched for those associated with extracellular organization, caspase activity, MAPK signaling, and eicosanoid metabolism; the latter two terms are also among the top pathways in our multi-omic analyses consistent with widespread changes across transcriptional, epigenetic, and topological levels. Prostaglandins analogues are widely used as the first line of treatment for POAG; reducing IOP by increasing the AH outflow through conventional and unconventional pathways (Angeli and Supuran, 2019; Tejwani et al., 2020; Winkler and Fautsch, 2014). Pathways that gain SEs post treatment also identify apoptotic signaling, suggesting a potential role of the gain of CEBPD binding in regulating pathways associated with increased IOP.

Genes predicted to have reduced JUN binding are enriched for integrin function, adherens junction organization, and cell surface receptor signaling pathway. Focal adhesions which involve integrins remodel the extracellular matrix, resulting in cross-linked actin network (CLANs) formation in glaucoma (Filla et al., 2017). Integrins interact with receptors for advanced glycation end-products (RAGEs) to regulate the level of glycosylated proteins in podocytes (Kim and Dryer, 2021). AGE-RAGE signaling was one of the top pathways identified by our multi-omic analysis in dexamethasone treated TM cells. AGEs are the result of high glucose, oxidative stress and advanced age; all of these have been implicated in vision loss, where they mediate crosslinking of extracellular matrix proteins (Kandarakis et al., 2014). Single-cell RNA-seq of TM derived from aging macaque eyes have ascertained a cluster enriched for AGE-RAGE signaling pathway (Wu et al., 2024). These results therefore direct to association of AGE-RAGE to glaucomatous conditions.

### The genomic context of POAG and IOP SNPs

GWAS studies have discovered genetic loci linked to POAG and IOP. Our study builds on these findings by uncovering specific target genes for disease-associated loci. More than half all promoter-SNP loops we identified are not observed in any of 20 tissues previously examined by promoter-capture Hi-C, thereby exposing novel targets. We have identified *MIR760* gene as the target gene for *BCAR3* locus and the lead variant rs7525880 that are associated with POAG (Han *et al*., 2023). *MIR760* is downregulated in human primary TM cells treated with either TGFβ1 or TGFβ2 (Doyle et al., 2024). Additionally, *MIR760* is downregulated in glaucoma tissues, when compared with their expression levels in controls (Zhao et al., 2019). We have also identified *FZD4* gene as the candidate target for POAG GWAS locus *TMEM135* and the lead variant rs2513214 (Han *et al*., 2023). FZD4 belongs to frizzled receptors family and its stimulation can activate both Wnt/β-catenin canonical and Wnt/Ca²⁺ non canonical pathways (Shyam et al., 2010). *FZD4* is upregulated in TM cells subjected to cyclic mechanical stretch as compared to control cells (Soundararajan et al., 2022). We have shown that Wnt signaling pathway is broadly enriched for epigenetic changes following dexamethasone treatment in TM cells. TGFβ and Wnt signaling pathways are reported to be crucial for maintaining TM homeostasis and regulating IOP (Fuchshofer and Tamm, 2012; Webber et al., 2018).

SEs harboring POAG lead variants are also rarely detected in non-TM samples. We have connected multiple SNPs residing in a SE to *JAG1*, which encodes a canonical ligand for NOTCH receptor that mediates downstream functions in development, homeostasis and repair. Intriguingly, *JAG1* plays a key role in kidney fibrosis (Huang et al., 2018), a condition similar to the fibrosis observed in glaucoma (McDonnell et al., 2014). Additionally, the expression of *JAG1* and Notch receptors progressively decline with increasing stiffness of TM cells, a condition that mimics glaucoma (Dhamodaran et al., 2022). We hypothesize that the variation in *JAG1* regulation at this SE may contribute to IOP changes in TM cells and therefore to glaucoma risk.

In sum, our high-resolution promoter-centered chromatin contact maps and epigenetic signatures have provided unique insights into TM-specific gene regulation. Through non-coding genome annotation and physical contact mapping, we have also uncovered new genes associated with elevated IOP and POAG pathogenesis. We however appreciate that experimental validations are required for the new target genes identified here. Further investigations would be necessary to examine 3-D chromatin architecture of donor TM tissue from controls and glaucoma patients. Owing to heterogeneity in cellular composition of TM, single-cell studies including single cell Hi-C will help in delineating cell type specificity of chromatin contacts.

## Limitations of the study

While our study maps the first high resolution genome topology map of control and dexamethasone treated TM cells. Further validations are required for the new target genes identified in the study. The study lacks comparison of chromatin map and epigenetic signatures to control TM tissue and glaucomatous cells. Owing to heterogeneity in cellular composition of TM performing single-cell Hi-C will define cell type specificity of chromatin contacts.

## RESOURCE AVAILABILITY

### Lead contact

Further information and requests for resources and reagents should be directed to and will be fulfilled by the lead contact, Anand Swaroop.

### Materials availability

All unique/stable reagents generated in this study are available with a completed Materials Transfer Agreement per NIH policy. Further information and requests for resources and materials should be directed to and will be fulfilled by the lead contact, Anand Swaroop.

### Data and code availability

- All datasets used from public resources or produced in this study are summarized in the key resources table. The next generation sequencing data generated in this study are available at the Gene Expression Omnibus (GEO; GSE301525).

- To review GEO accession GSE301525: Go to https://www.ncbi.nlm.nih.gov/geo/query/acc.cgi?acc=GSE301525 Enter token gdmrwcqizvebtqn into the box
- Any additional information required to reanalyze the data reported in this paper is available from the lead contact upon request.

## ACKNOWLEDGMENTS

We thank members of Swaroop laboratory, especially Claire Marchal, for discussions, Dr. Saidas Nair for his expertise in glaucoma genetics and Dr. Greg Germino for sharing unpublished insights. This work was supported by funding from the National Eye Institute (ZIAEY000450, ZIAEY000546, and R01EY036373) and utilized the computational resources of the NIH HPC Biowulf cluster (https://hpc.nih.gov). The contributions of the NIH authors are considered Works of the United States Government. The findings and conclusions presented in this paper are those of the authors and do not necessarily reflect the views of the NIH or the U.S. Department of Health and Human Services.

## AUTHOR CONTRIBUTIONS

Conceptualization, N.S. and A.S.; Methodology and Investigation, N.S., M.A.E., R.M., P.V.R.; Bioinformatic analysis and Visualization, Z.B., J.A.; Data submission, Z.B.; Writing - Original Draft, N.S., Z.B., J.A., A.S.; Writing, Review & Editing, all authors; Supervision, Project administration, and Funding acquisition, V.R., A.S.

## MATERIALS AND METHODS

### Key Resources

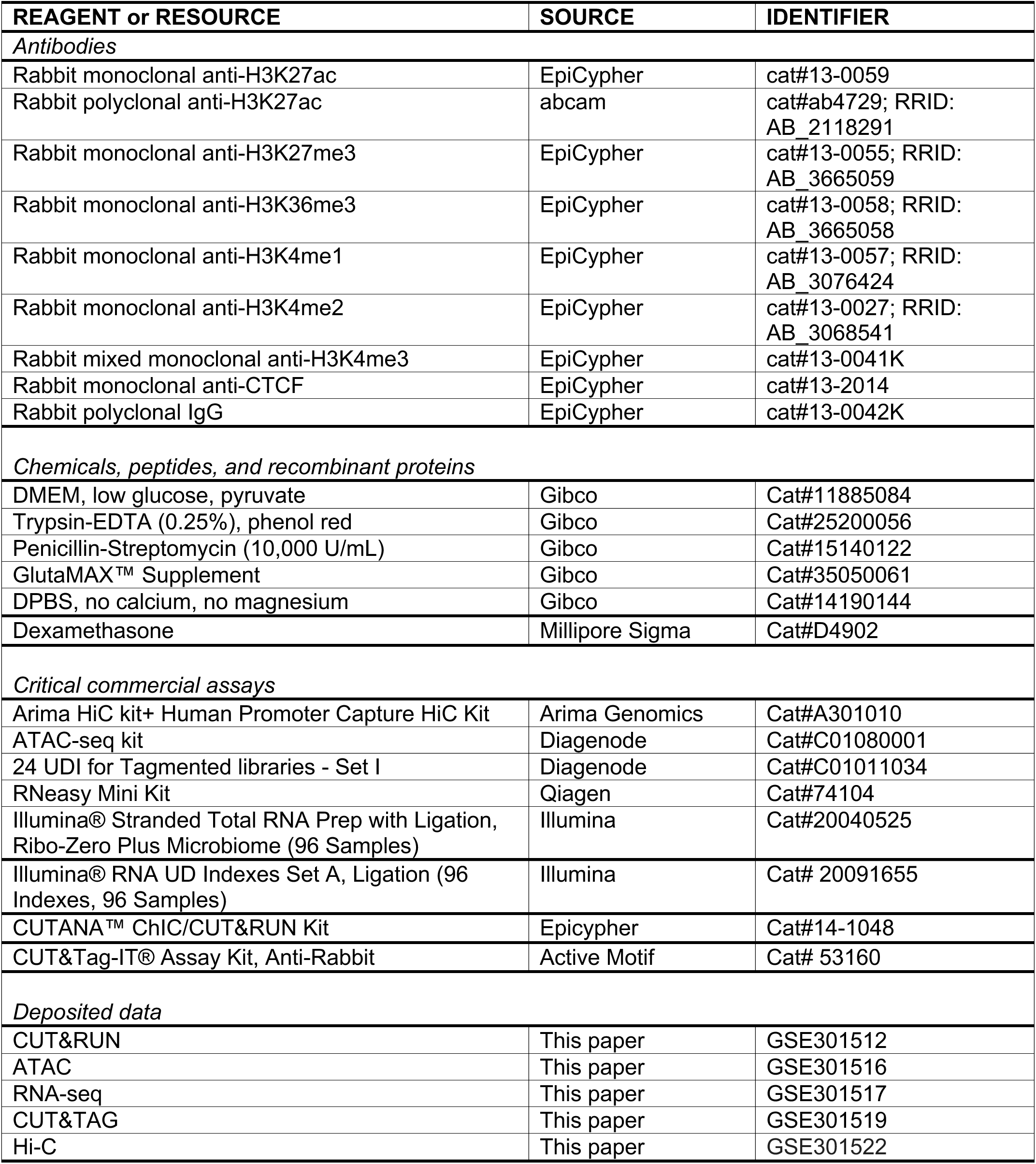
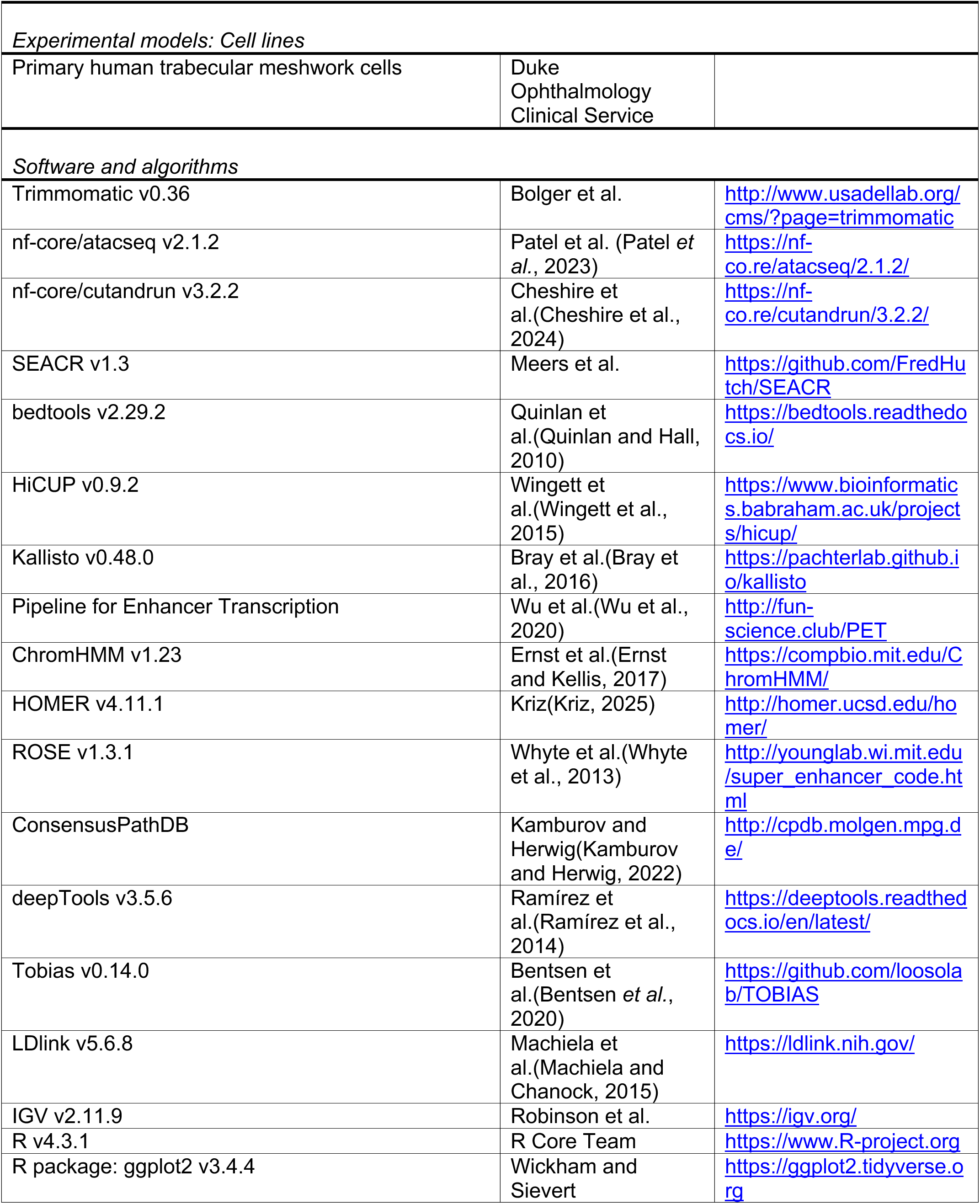
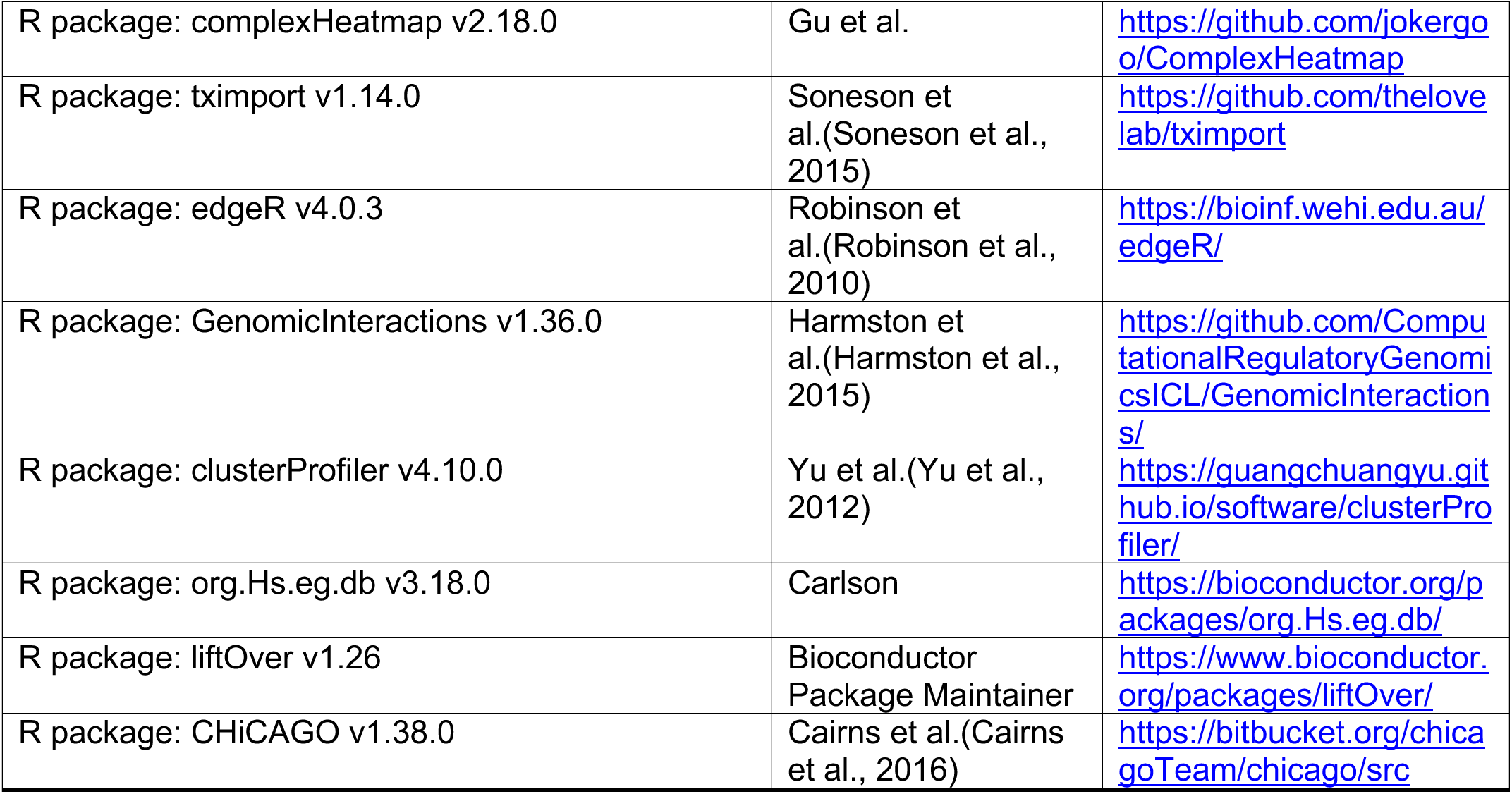

## Experimental Methods

### Human Trabecular Meshwork Cells (TMCs) Culture

Trabecular meshwork primary cells were cultured from human TM tissue derived from healthy donor corneal rings (ages 62, 70, 72, 76 and 83 years), leftover from corneal transplantation surgeries performed at the Duke Ophthalmology Clinical Service. TM cells isolation, characterization and culture was performed as per consensus recommendations for TM cells culture(Keller et al., 2018). Primary TM cells were cultured in Dulbecco’s Modified Eagle complete growth medium (DMEM) containing low glucose (cat#11885084) with 10% heat inactivated FBS (fetal bovine serum), 1X Penicillin-Streptomycin (cat#15140122) and GlutaMAX supplement (cat#35050061) at 37°C in an aseptic incubator at 5% CO2. All experiments on primary TM cells were performed between passage 2-6.

TM cells with 80% confluency were maintained in medium containing 2% FBS for 2 hours before treating with 0.5 μM dexamethasone (cat#D4902) and ethanol as a vehicle control. The cells were treated for 7 days with media containing 2% FBS and dexamethasone changed every other day.

### Total RNA-seq

We performed total RNA-sequencing on three samples each for the resource, dexamethasone control, and dexamethasone treatment experiments. For each experiment, primary TM cells were trypsinized, flash frozen and kept at −80°C. Total RNA was isolated using Qiagen RNeasy Mini Kit (Germantown, MD, USA; cat# 74104) and the quality of RNA was determined using Agilent TapeStation (Santa Clara, CA, USA). Libraries were prepared from 10ng of total RNA using the Illumina Stranded Total RNA Prep, Ligation with Ribo-Zero Plus kit (San Diego, CA, USA; cat# 20040525). Libraries were sequenced on an Illumina NextSeq 2000 platform and reads were trimmed to remove low quality bases using Trimmomatic v0.36(Bolger et al., 2014).

### Bulk Assay for Transposase-Accessible Chromatin using sequencing (ATAC-seq)

We performed ATAC-seq on two samples each for the resource, control, and dexamethasone treatment experiment. For each, approximately 50,000 cells were used for cell lysis, nuclei extraction, tagmentation and DNA purification. ATAC-seq and library preparation was performed using a commercial kit from Diagenode (cat#C01080001, cat# C01011034) following the manufacturer’s instructions. Pair-end sequencing with read length of 50 base pairs was done on the NextSeq 2000 platform (Illumina, CA, USA). Reads were processed using the nf-core ATACseq pipeline (Patel *et al*., 2023) using the default settings with peaks called using SEACR v1.3 (Meers et al., 2019). We retained all replicate peaks identified in the conservative set by irreproducible discovery rate (IDR) analysis.

### Cleavage under targets and release using nuclease (CUT&RUN)

The samples were prepared from primary cultures of 200,000-300,000 TM cells of three different individuals for CTCF (cat#13-2014), IgG (cat#13-0042K), and each of the following histone marks: H3K27ac (cat#13-0059, cat#ab4729), H3K27me3 (cat#13-0055), H3K4me1 (cat#13-0057), H3K4me2 (cat#13-0027), H3K4me3 (cat#13-0041K), H3K36me3 (cat#13-0058), and H3K9me3 (cat# ab8898). Sample libraries were prepared using the CUTANA ChiC/CUT&RUN kit (cat#14-1048) and sequenced on the NextSeq 2000 platform (Illumina, CA, USA) at a read length of 50 base pairs. Reads were processed using the nfcore cutandrun pipeline v3.2.2 (Cheshire *et al*., 2024). Peaks were called using SEACR v1.3(Meers *et al*., 2019) and overlapped using bedtools v2.29.2 (Quinlan and Hall, 2010); only peaks observed in at least two samples were retained for further analysis.

### Cleavage Under Targets and Tagmentation (CUT&Tag)

Two biological replicates for the control and dexamethasone treated experiment were used for CUT&Tag to detect H3K36me3, H3K4me1, H3K4me2, H3K4me3, and CTCF. Three biological replicates were used for H3K27ac and H3K27me3. CUT&Tag experiment was performed on 100,000-200,000 cells using CUT&Tag-IT® Assay Kit, Anti-Rabbit (cat# 53171). The libraries were sequenced on the NextSeq 2000 platform (Illumina, CA, USA) at a read length of 50 base pairs. CUT&Tag samples were processed using the nfcore cutandrun pipeline (Cheshire *et al*., 2024) with peaks called using SEACR v1.3 (Meers *et al*., 2019); for each mark, peaks called in at least two samples were retained for downstream analyses.

### Promoter Capture Hi-C

For each experiment, approximately five million cells from three independent TM primary cell strains were used for capture Hi-C. Cells were crosslinked with 2% formaldehyde in PBS for 10 mins, quenched with 125mM glycine for 5 min at room temperature, lysed and processed using Arima-HiC+ kit, the Arima Human Promoter Panel, and the Arima Library Prep Module (cat#A301010, Arima Genomics, CA, USA) according to the manufacturer’s protocols. Each library was sequenced twice, once before and once after target capture, on the NextSeq 2000 platform (Illumina, CA, USA) at a read length of 150 base pairs. Hi-C reads were filtered and mapped via HiCUP v0.9.2 (Wingett *et al*., 2015) using the *in silico* digested hg38 genome provided by Arima.

### Transcriptome Analysis

Transcript level quantitation was performed using Kallisto v0.48.0 (Bray *et al*., 2016) employing a merged transcript cDNA and ncRNA FASTA file downloaded from Ensembl v106. Gene level quantification was generated by summarization of the transcript level quantitation using tximport v1.14.0 (Soneson *et al*., 2015) with using the lengthScaledTPM option. The gene level count values were TMM normalized and differential expression in the dexamethasone experiment was performed using edgeR v4.0.3 (Robinson *et al*., 2010). Enrichment of GO terms and KEGG pathways was performed using clusterProfiler v4.10.0 (Yu *et al*., 2012).

### Enhancer RNA identification and analysis

We predicted eRNA from total RNA as described previously (Zhu et al., 2013). In brief, enhancer regions identified from ChromHMM were merged within 50bp. Next, we removed regions that overlapped gene bodies, located within 3kb of a TSS, or located within 3kb of an H3K4me3 peak. The remaining set of 30,416 putative eRNAs were used as a reference to quantify expression using the Pipeline for Enhancer Transcription (Wu *et al*., 2020). We retained 4,152 putative eRNA with >1 CPM expression in at least 2 samples as the final set.

### Annotation of Cis-regulatory elements, enhancers, and super enhancers

Chromatin was annotation was performed by integrating histone marks and CTCF peaks from CUT&RUN with chromatin accessibility data following the technique described previously (Marchal et al., 2022). Briefly, we transformed read coverage into binaries in 500bp windows then computed models with 5 to 20 chromatin states using ChromHMM v1.23 (Ernst and Kellis, 2017). For each model, we calculated the mean correlation between each chromatin state and the most similar chromatin state in each model with additional states. This correlation plateaued at 13 states in the resource experiment and 10 states in the dexamethasone experiment indicating that additional chromatin states provide minimal information. For each state of the final state models, the average coverage for the chromatin marks and accessibility were computed using HOMER v4.11.1 (Kriz, 2025) and the chromatin signatures for each segment were used to manually infer a biological annotation of each state. All states annotated as promoter or enhancer were merged to create a set of CREs. Enhancer states were further annotated based on their histone marks as described previously (Barral and Dejardin, 2023).

Next, CREs were integrated with H3K27ac coverage to identify super enhancers using ROSE v1.3.1 (Whyte *et al*., 2013). For the resource set, SEs were identified by integration with six sets of H3K27ac coverage generated using two sets of antibodies from three biological replicates; SE present in at least 4 sets were retained. For the dexamethasone experiment, three sets of H3K27ac reads were used for each condition and SEs present in at least two replicates were retained. SEs lost or gained following dexamethasone treatment were identified by overlapping the control and treated SEs using intersect from bedtools v2.29.2 (Quinlan and Hall, 2010).

Expressed genes overlapping SEs of interest were to perform gene set enrichment analyses using clusterProfiler 4.10.0 (Yu *et al*., 2012) and ConsensusPathDB (Kamburov and Herwig, 2022). To determine tissue specificity, we used bedtools intersect to count the number of samples from SEdb 2.0 (Wang *et al*., 2023) with an SE overlapping at least 70% of a TM SE. For visualization, we generated a bedgraph file counting the number of samples overlapping each 50bp window across the genome.

### Promoter capture Hi-C Analysis

Compartments were called from pre-capture libraries with HOMER v4.11.1 (Kriz, 2025) using 100 kb sliding windows with a step of 50 kb (via the superRes and res options, respectively). To find compartment changes after dexamethasone treatment, .hic files were converted to sparse matrices using strawr v0.0.92 (Durand et al., 2016) then tested using dcHiC (Chakraborty et al., 2022). Changes in histone marks and accessibility within dynamic compartments were identified by first computing the average change in signal for each mark across each pair of treated and untreated samples using bamCompare from deepTools v3.5.6 (Ramírez *et al*., 2014). Next, we filtered to 50kb windows that significantly shifted towards A or B compartment and tested if the mean change in signal for these regions differed from zero using a one t-test.

Loops were identified from post-capture libraries using CHiCAGO (Cairns *et al*., 2016) via the Arima CHiC pipeline. Only loops present at least 2 samples were retained for further analysis. In the dexamethasone experiment, loops were identified as stable, lost, or gained based on two-anchor overlap using GenomicInteractions v1.36.0 (Harmston *et al*., 2015). Similarly, loops contacting promoters, enhancers, and repressors were identified using GenomicInteractions. Gene promoters for transcription factors were derived from previously published data (Lambert et al., 2018).

To compare TM promoter-centered loops in the resource experiment to existing pcHiC data (Jung *et al*., 2019), we used liftOver v1.26 in R to convert TM loops to hg19; we retained only loops where both anchors could be converted successfully. Next, loops were filtered to include only those that overlapped at least one targeted promoter in the previous pcHiC study. In total, 29,468 (97.0%) of loops were retained were retained through this process. Finally, loops overlap was compared using GenomicInteractions v1.36.0. All contact maps were created by converting filtered bam files to hic files using HOMER v4.11.1 then visualized in Juicebox.

### Chromatin accessibility and ATAC footprinting

Changes in chromatin accessibility were identified by overlapping control and dexamethasone treated peak lists using bedtools intersect. To assess differential TF binding, bam files from control and dexamethasone-treated TM samples were used as input for Tobias v0.14.0 (Bentsen *et al*., 2020) implemented via the CCBR pipeline (Koparde and Sovacool, 2025). TF binding was predicted based using the position frequency matrices from JASPAR core, non-redundant database (Rauluseviciute et al., 2024).

### Identification of linkage disequilibrium (LD) variants for IOP and POAG GWAS loci

Lead variants from two POAG (Gharahkhani *et al*., 2021; Han *et al*., 2023) and three IOP (Gharahkhani *et al*., 2021; Khawaja *et al*., 2018; MacGregor *et al*., 2018) GWAS studies were included for the analysis. We calculated linkage disequilibrium (LD) for all lead variants within 1 MB region (Gharahkhani *et al*., 2021; Han *et al*., 2023; Khawaja *et al*., 2018; MacGregor *et al*., 2018) using LDlink v5.6.8 suite(Machiela and Chanock, 2015) and LDproxy module using individuals with European ancestry from 1000 Genomes Project data (Genomes Project *et al*., 2015).

### Integration of promoter loops and SEs for identification of target genes for IOP and POAG GWAS loci

To identify target genes for variants identified in IOP and POAG GWAS studies (Gharahkhani *et al*., 2021; Han *et al*., 2023; Khawaja *et al*., 2018; MacGregor *et al*., 2018), we overlapped the variants with promoter loops and SEs. Identification of target gene for variants forming chromatin loop, one foot of the loop overlaps the lead or LD variant, and the second foot of the loop overlaps the promoter of a gene. SE target genes were defined by the lead or LD variant and promoter of the gene overlapping the same SE. The closest target genes overlapping with loops and SEs for lead or LD variants were obtained using the closestBed command from bedtools v2.31.1 (Quinlan and Hall, 2010).

### Integrated Analysis

We identified genes undergoing a variety of changes across chromatin topology, epigenetic mechanism, and expression after dexamethasone treatment. We tested for pathway enrichment among 16 gene sets. Specifically, we examined differentially expressed genes and genes within changing chromatin compartments. We also tested genes with changes in chromatin accessibility and overlapping each gained or lost SEs, CREs, activated enhancers as well as changes in each of the following epigenetic marks: H3K27ac, H3K27me3, H3K4me1, H3K4me3, H3K36me3, and CTCF. Finally, we examined altered looping including the genes with the greatest number of loop changes (top 5%) and gained or lost loops between promoters and enhancers, repressors, or SEs. Each set was individually used for gene set enrichment against GO and KEGG terms using ClusterProfiler (Yu *et al*., 2012); terms appearing in at least 4 sets were plotted using ComplexHeatmap.

## REFERENCES

Allison, K., Patel, D., and Alabi, O. (2020). Epidemiology of Glaucoma: The Past, Present, and Predictions for the Future. Cureus 12, e11686. 10.7759/cureus.11686.

Angeli, A., and Supuran, C.T. (2019). Prostaglandin receptor agonists as antiglaucoma agents (a patent review 2013 - 2018). Expert Opin Ther Pat 29, 793–803. 10.1080/13543776.2019.1661992.

Banner, L.R., Patterson, P.H., Allchorne, A., Poole, S., and Woolf, C.J. (1998). Leukemia inhibitory factor is an anti-inflammatory and analgesic cytokine. J Neurosci 18, 5456–5462. 10.1523/JNEUROSCI.18-14-05456.1998.

Barral, A., and Dejardin, J. (2023). The chromatin signatures of enhancers and their dynamic regulation. Nucleus 14, 2160551. 10.1080/19491034.2022.2160551.

Baudouin, C., Kolko, M., Melik-Parsadaniantz, S., and Messmer, E.M. (2021). Inflammation in Glaucoma: From the back to the front of the eye, and beyond. Prog Retin Eye Res 83, 100916. 10.1016/j.preteyeres.2020.100916.

Bentsen, M., Goymann, P., Schultheis, H., Klee, K., Petrova, A., Wiegandt, R., Fust, A., Preussner, J., Kuenne, C., Braun, T., et al. (2020). ATAC-seq footprinting unravels kinetics of transcription factor binding during zygotic genome activation. Nat Commun 11, 4267. 10.1038/s41467-020-18035-1.

Bermudez, J.Y., Webber, H.C., Brown, B., Braun, T.A., Clark, A.F., and Mao, W. (2017). A Comparison of Gene Expression Profiles between Glucocorticoid Responder and Non-Responder Bovine Trabecular Meshwork Cells Using RNA Sequencing. PLoS One 12, e0169671. 10.1371/journal.pone.0169671.

Binder, E.B. (2009). The role of FKBP5, a co-chaperone of the glucocorticoid receptor in the pathogenesis and therapy of affective and anxiety disorders. Psychoneuroendocrinology 34 *Suppl 1*, S186–195. 10.1016/j.psyneuen.2009.05.021.

Bolger, A.M., Lohse, M., and Usadel, B. (2014). Trimmomatic: a flexible trimmer for Illumina sequence data. Bioinformatics 30, 2114–2120. 10.1093/bioinformatics/btu170.

Bray, N.L., Pimentel, H., Melsted, P., and Pachter, L. (2016). Near-optimal probabilistic RNA-seq quantification. Nat Biotechnol 34, 525–527. 10.1038/nbt.3519.

Bresson, E., Boucher-Kovalik, S., Chapdelaine, P., Madore, E., Harvey, N., Laberge, P.Y., Leboeuf, M., and Fortier, M.A. (2011). The human aldose reductase AKR1B1 qualifies as the primary prostaglandin F synthase in the endometrium. J Clin Endocrinol Metab 96, 210–219. 10.1210/jc.2010-1589.

Buenrostro, J.D., Wu, B., Chang, H.Y., and Greenleaf, W.J. (2015). ATAC-seq: A Method for Assaying Chromatin Accessibility Genome-Wide. Curr Protoc Mol Biol 109, 21 29 21-21 29 29. 10.1002/0471142727.mb2129s109.

Burghardt, T., Kastner, J., Suleiman, H., Rivera-Milla, E., Stepanova, N., Lottaz, C., Kubitza, M., Boger, C.A., Schmidt, S., Gorski, M., et al. (2013). LMX1B is essential for the maintenance of differentiated podocytes in adult kidneys. J Am Soc Nephrol 24, 1830–1848. 10.1681/ASN.2012080788.

Cairns, J., Freire-Pritchett, P., Wingett, S.W., Várnai, C., Dimond, A., Plagnol, V., Zerbino, D., Schoenfelder, S., Javierre, B.M., Osborne, C., et al. (2016). CHiCAGO: robust detection of DNA looping interactions in Capture Hi-C data. Genome Biol 17. ARTN 127 10.1186/s13059-016-0992-2.

Cela, D., Brignole-Baudouin, F., Labbe, A., and Baudouin, C. (2021). The trabecular meshwork in glaucoma: An inflammatory trabeculopathy? J Fr Ophtalmol 44, e497–e517. 10.1016/j.jfo.2021.09.001.

Chakraborty, A., Wang, J.G., and Ay, F. (2022). dcHiC detects differential compartments across multiple Hi-C datasets. Nat Commun 13, 6827. 10.1038/s41467-022-34626-6.

Cheshire, C., West, C., Rönkkö, T., Patel, H., Hodgetts, T., Ladd, D., Thiery, A., Fields, C., Deu-Pons, J., Ewels, P., et al. (2024). nf-core/cutandrun: nf-core/cutandrun v3.2.2. 10.5281/zenodo.5653535.

Choi, D., Kang, W., Park, S., Son, B., and Park, T. (2023). Identification of Glucocorticoid Receptor Target Genes That Potentially Inhibit Collagen Synthesis in Human Dermal Fibroblasts. Biomolecules 13. 10.3390/biom13060978.

Clark, J.D., Lin, L.L., Kriz, R.W., Ramesha, C.S., Sultzman, L.A., Lin, A.Y., Milona, N., and Knopf, J.L. (1991). A novel arachidonic acid-selective cytosolic PLA2 contains a Ca(2+)-dependent translocation domain with homology to PKC and GAP. Cell 65, 1043–1051. 10.1016/0092-8674(91)90556-e.

Conlon, R., Saheb, H., and Ahmed, II (2017). Glaucoma treatment trends: a review. Can J Ophthalmol 52, 114–124. 10.1016/j.jcjo.2016.07.013.

Craig, J.E., Han, X., Qassim, A., Hassall, M., Cooke Bailey, J.N., Kinzy, T.G., Khawaja, A.P., An, J., Marshall, H., Gharahkhani, P., et al. (2020). Multitrait analysis of glaucoma identifies new risk loci and enables polygenic prediction of disease susceptibility and progression. Nat Genet 52, 160–166. 10.1038/s41588-019-0556-y.

Cross, S.H., Macalinao, D.G., McKie, L., Rose, L., Kearney, A.L., Rainger, J., Thaung, C., Keighren, M., Jadeja, S., West, K., et al. (2014). A dominant-negative mutation of mouse Lmx1b causes glaucoma and is semi-lethal via LDB1-mediated dimerization [corrected]. PLoS genetics 10, e1004359. 10.1371/journal.pgen.1004359.

Dehingia, B., Milewska, M., Janowski, M., and Pekowska, A. (2022). CTCF shapes chromatin structure and gene expression in health and disease. EMBO Rep 23, e55146. 10.15252/embr.202255146.

Dhamodaran, K., Baidouri, H., Nartey, A., Staverosky, J., Keller, K., Acott, T., Vranka, J.A., and Raghunathan, V.K. (2022). Endogenous expression of Notch pathway molecules in human trabecular meshwork cells. Exp Eye Res 216, 108935. 10.1016/j.exer.2022.108935.

Doyle, C., Callaghan, B., Roodnat, A.W., Armstrong, L., Lester, K., Simpson, D.A., Atkinson, S.D., Sheridan, C., McKenna, D.J., and Willoughby, C.E. (2024). The TGFbeta Induced MicroRNAome of the Trabecular Meshwork. Cells 13. 10.3390/cells13121060.

Dreyer, S.D., Zhou, G., Baldini, A., Winterpacht, A., Zabel, B., Cole, W., Johnson, R.L., and Lee, B. (1998). Mutations in LMX1B cause abnormal skeletal patterning and renal dysplasia in nail patella syndrome. Nat Genet 19, 47–50. 10.1038/ng0598-47.

Durand, N.C., Robinson, J.T., Shamim, M.S., Machol, I., Mesirov, J.P., Lander, E.S., and Aiden, E.L. (2016). Juicebox Provides a Visualization System for Hi-C Contact Maps with Unlimited Zoom. Cell Syst 3, 99–101. 10.1016/j.cels.2015.07.012.

Ernst, J., and Kellis, M. (2017). Chromatin-state discovery and genome annotation with ChromHMM. Nat Protoc 12, 2478–2492. 10.1038/nprot.2017.124.

Fan, B.J., Wang, D.Y., Tham, C.C., Lam, D.S., and Pang, C.P. (2008). Gene expression profiles of human trabecular meshwork cells induced by triamcinolone and dexamethasone. Invest Ophthalmol Vis Sci 49, 1886–1897. 10.1167/iovs.07-0414.

Farley, F.A., Lichter, P.R., Downs, C.A., McIntosh, I., Vollrath, D., and Richards, J.E. (1999). An orthopaedic scoring system for nail-patella syndrome and application to a kindred with variable expressivity and glaucoma. J Pediatr Orthop 19, 624–631.

Filla, M.S., Faralli, J.A., Peotter, J.L., and Peters, D.M. (2017). The role of integrins in glaucoma. Exp Eye Res 158, 124–136. 10.1016/j.exer.2016.05.011.

Friedman, D.S., Wolfs, R.C., O’Colmain, B.J., Klein, B.E., Taylor, H.R., West, S., Leske, M.C., Mitchell, P., Congdon, N., Kempen, J., and Eye Diseases Prevalence Research, G. (2004). Prevalence of open-angle glaucoma among adults in the United States. Arch Ophthalmol 122, 532–538. 10.1001/archopht.122.4.532.

Fuchshofer, R., and Tamm, E.R. (2012). The role of TGF-beta in the pathogenesis of primary open-angle glaucoma. Cell Tissue Res 347, 279–290. 10.1007/s00441-011-1274-7.

Garg, A., and Gazzard, G. (2020). Treatment choices for newly diagnosed primary open angle and ocular hypertension patients. Eye (Lond) 34, 60–71. 10.1038/s41433-019-0633-6.

Garufi, A., Pistritto, G., and D’Orazi, G. (2023). HIPK2 as a Novel Regulator of Fibrosis. Cancers (Basel) 15. 10.3390/cancers15041059.

Genomes Project, C., Auton, A., Brooks, L.D., Durbin, R.M., Garrison, E.P., Kang, H.M., Korbel, J.O., Marchini, J.L., McCarthy, S., McVean, G.A., and Abecasis, G.R. (2015). A global reference for human genetic variation. Nature 526, 68–74. 10.1038/nature15393.

Gharahkhani, P., Jorgenson, E., Hysi, P., Khawaja, A.P., Pendergrass, S., Han, X., Ong, J.S., Hewitt, A.W., Segre, A.V., Rouhana, J.M., et al. (2021). Genome-wide meta-analysis identifies 127 open-angle glaucoma loci with consistent effect across ancestries. Nat Commun 12, 1258. 10.1038/s41467-020-20851-4.

Gramer, G., Weber, B.H., and Gramer, E. (2014). Results of a patient-directed survey on frequency of family history of glaucoma in 2170 patients. Invest Ophthalmol Vis Sci 55, 259–264. 10.1167/iovs.13-13020.

Hamel, A.R., Yan, W., Rouhana, J.M., Monovarfeshani, A., Jiang, X., Mehta, P.A., Advani, J., Luo, Y., Liang, Q., Rajasundaram, S., et al. (2024). Integrating genetic regulation and single-cell expression with GWAS prioritizes causal genes and cell types for glaucoma. Nat Commun 15, 396. 10.1038/s41467-023-44380-y.

Han, X., Gharahkhani, P., Hamel, A.R., Ong, J.S., Renteria, M.E., Mehta, P., Dong, X., Pasutto, F., Hammond, C., Young, T.L., et al. (2023). Large-scale multitrait genome-wide association analyses identify hundreds of glaucoma risk loci. Nat Genet 55, 1116–1125. 10.1038/s41588-023-01428-5.

Harmston, N., Ing-Simmons, E., Perry, M., Baresic, A., and Lenhard, B. (2015). GenomicInteractions: An R/Bioconductor package for manipulating and investigating chromatin interaction data. Bmc Genomics 16. ARTN 963 10.1186/s12864-015-2140-x.

Hershko, A., and Ciechanover, A. (1998). The ubiquitin system. Annu Rev Biochem 67, 425–479. 10.1146/annurev.biochem.67.1.425.

Huang, S., Park, J., Qiu, C., Chung, K.W., Li, S.Y., Sirin, Y., Han, S.H., Taylor, V., Zimber-Strobl, U., and Susztak, K. (2018). Jagged1/Notch2 controls kidney fibrosis via Tfam-mediated metabolic reprogramming. PLoS Biol 16, e2005233. 10.1371/journal.pbio.2005233.

Ishimoto, Y., Menezes, L.F., Nakaya, N., Barbosa, K., Horie, Y., Yoshida, T., Reece, J., Zhou, F., Tomarev, S., Kerosuo, L., and Germino, G.G. (2026). Studies of mice with a large deletion of the ARPKD-associated Pkhd1 locus likely explain its GWAS association with glaucoma in humans. bioRxiv. 10.64898/2026.02.15.706040.

Jubb, A.W., Boyle, S., Hume, D.A., and Bickmore, W.A. (2017). Glucocorticoid Receptor Binding Induces Rapid and Prolonged Large-Scale Chromatin Decompaction at Multiple Target Loci. Cell Rep 21, 3022–3031. 10.1016/j.celrep.2017.11.053.

Jung, I., Schmitt, A., Diao, Y., Lee, A.J., Liu, T., Yang, D., Tan, C., Eom, J., Chan, M., Chee, S., et al. (2019). A compendium of promoter-centered long-range chromatin interactions in the human genome. Nat Genet 51, 1442–1449. 10.1038/s41588-019-0494-8.

Kamburov, A., and Herwig, R. (2022). ConsensusPathDB 2022: molecular interactions update as a resource for network biology. Nucleic Acids Res 50, D587–D595. 10.1093/nar/gkab1128.

Kandarakis, S.A., Piperi, C., Topouzis, F., and Papavassiliou, A.G. (2014). Emerging role of advanced glycation-end products (AGEs) in the pathobiology of eye diseases. Prog Retin Eye Res 42, 85–102. 10.1016/j.preteyeres.2014.05.002.

Kathirvel, K., Lester, K., Haribalaganesh, R., Krishnadas, R., Muthukkaruppan, V., Lane, B., Simpson, D.A., Goljanek-Whysall, K., Sheridan, C., Bharanidharan, D., et al. (2022). Short and long-term effect of dexamethasone on the transcriptome profile of primary human trabecular meshwork cells in vitro. Sci Rep 12, 8299. 10.1038/s41598-022-12443-7.

Keller, K.E., Bhattacharya, S.K., Borras, T., Brunner, T.M., Chansangpetch, S., Clark, A.F., Dismuke, W.M., Du, Y., Elliott, M.H., Ethier, C.R., et al. (2018). Consensus recommendations for trabecular meshwork cell isolation, characterization and culture. Exp Eye Res 171, 164–173. 10.1016/j.exer.2018.03.001.

Khawaja, A.P., Cooke Bailey, J.N., Wareham, N.J., Scott, R.A., Simcoe, M., Igo, R.P., Jr., Song, Y.E., Wojciechowski, R., Cheng, C.Y., Khaw, P.T., et al. (2018). Genome-wide analyses identify 68 new loci associated with intraocular pressure and improve risk prediction for primary open-angle glaucoma. Nat Genet 50, 778–782. 10.1038/s41588-018-0126-8.

Kim, E.Y., and Dryer, S.E. (2021). RAGE and alphaVbeta3-integrin are essential for suPAR signaling in podocytes. Biochim Biophys Acta Mol Basis Dis 1867, 166186. 10.1016/j.bbadis.2021.166186.

Kinoshita, M., Nakamura, T., Ihara, M., Haraguchi, T., Hiraoka, Y., Tashiro, K., and Noda, M. (2001). Identification of human endomucin-1 and −2 as membrane-bound O-sialoglycoproteins with anti-adhesive activity. FEBS Lett 499, 121–126. 10.1016/s0014-5793(01)02520-0.

Koparde, V., and Sovacool, K. (2025). CCBR TOBIAS snakemake pipeline (v0.3.1). https://zenodo.org/records/15722020.

Kriz, A.J. (2025). HOMER for Analysis of Hi-C Data and Assessment of Composite Structure of the X Chromosome. Methods Mol Biol 2856, 71–78. 10.1007/978-1-0716-4136-1_5.

Lambert, S.A., Jolma, A., Campitelli, L.F., Das, P.K., Yin, Y., Albu, M., Chen, X., Taipale, J., Hughes, T.R., and Weirauch, M.T. (2018). The Human Transcription Factors. Cell 172, 650–665. 10.1016/j.cell.2018.01.029.

Li, L., Huang, J., and Liu, Y. (2023). The extracellular matrix glycoprotein fibrillin-1 in health and disease. Front Cell Dev Biol 11, 1302285. 10.3389/fcell.2023.1302285.

Liu, C., Shao, Z.M., Zhang, L., Beatty, P., Sartippour, M., Lane, T., Livingston, E., and Nguyen, M. (2001). Human endomucin is an endothelial marker. Biochem Biophys Res Commun 288, 129–136. 10.1006/bbrc.2001.5737.

Liu, P., and Johnson, R.L. (2010). Lmx1b is required for murine trabecular meshwork formation and for maintenance of corneal transparency. Dev Dyn 239, 2161–2171. 10.1002/dvdy.22347.

MacGregor, S., Ong, J.S., An, J., Han, X., Zhou, T., Siggs, O.M., Law, M.H., Souzeau, E., Sharma, S., Lynn, D.J., et al. (2018). Genome-wide association study of intraocular pressure uncovers new pathways to glaucoma. Nat Genet 50, 1067–1071. 10.1038/s41588-018-0176-y.

Machiela, M.J., and Chanock, S.J. (2015). LDlink: a web-based application for exploring population-specific haplotype structure and linking correlated alleles of possible functional variants. Bioinformatics 31, 3555–3557. 10.1093/bioinformatics/btv402.

Marchal, C., Singh, N., Batz, Z., Advani, J., Jaeger, C., Corso-Diaz, X., and Swaroop, A. (2022). High-resolution genome topology of human retina uncovers super enhancer-promoter interactions at tissue-specific and multifactorial disease loci. Nat Commun 13, 5827. 10.1038/s41467-022-33427-1.

Mathew, D.J., Maurya, S., Ho, J., Livne-Bar, I., Chan, D., Buys, Y., Sit, M., Trope, G., Flanagan, J.G., Gronert, K., and Sivak, J.M. (2025). Lipidomic Analysis Reveals Drug-Induced Lipoxins in Glaucoma Treatment. bioRxiv. 10.1101/2025.01.24.634771.

McDonnell, F., O’Brien, C., and Wallace, D. (2014). The role of epigenetics in the fibrotic processes associated with glaucoma. J Ophthalmol 2014, 750459. 10.1155/2014/750459.

McDonnell, F.S., Riddick, B.J., Roberts, H., Skiba, N., and Stamer, W.D. (2023). Comparison of the extracellular vesicle proteome between glaucoma and non-glaucoma trabecular meshwork cells. Front Ophthalmol (Lausanne) 3. 10.3389/fopht.2023.1257737.

Medina-Ortiz, W.E., Belmares, R., Neubauer, S., Wordinger, R.J., and Clark, A.F. (2013). Cellular fibronectin expression in human trabecular meshwork and induction by transforming growth factor-beta2. Invest Ophthalmol Vis Sci 54, 6779–6788. 10.1167/iovs.13-12298.

Meers, M.P., Tenenbaum, D., and Henikoff, S. (2019). Peak calling by Sparse Enrichment Analysis for CUT&RUN chromatin profiling. Epigenetics Chromatin 12, 42. 10.1186/s13072-019-0287-4.

Meng, H., Dai, Z., Zhang, W., Liu, Y., and Lai, L. (2018). Molecular mechanism of 15-lipoxygenase allosteric activation and inhibition. Phys Chem Chem Phys 20, 14785–14795. 10.1039/c7cp08586a.

Nott, A., Holtman, I.R., Coufal, N.G., Schlachetzki, J.C.M., Yu, M., Hu, R., Han, C.Z., Pena, M., Xiao, J., Wu, Y., et al. (2019). Brain cell type-specific enhancer-promoter interactome maps and disease-risk association. Science 366, 1134–1139. 10.1126/science.aay0793.

Ogryzko, V.V., Schiltz, R.L., Russanova, V., Howard, B.H., and Nakatani, Y. (1996). The transcriptional coactivators p300 and CBP are histone acetyltransferases. Cell 87, 953–959. 10.1016/s0092-8674(00)82001-2.

Oikonomidi, I., Burbridge, E., Cavadas, M., Sullivan, G., Collis, B., Naegele, H., Clancy, D., Brezinova, J., Hu, T., Bileck, A., et al. (2018). iTAP, a novel iRhom interactor, controls TNF secretion by policing the stability of iRhom/TACE. Elife 7. 10.7554/eLife.35032.

Overby, D.R., and Clark, A.F. (2015). Animal models of glucocorticoid-induced glaucoma. Exp Eye Res 141, 15–22. 10.1016/j.exer.2015.06.002.

Patel, H., Espinosa-Carrasco, J., Langer, B., Ewels, P., Garcia, M.U., Syme, R., Peltzer, A., Talbot, A., Behrens, D., Gabernet, G., et al. (2023). nf-core/atacseq: [2.1.2] - 2022-08-07 (2.1.2). 10.5281/zenodo.2634132.

Pattabiraman, P.P., Epstein, D.L., and Rao, P.V. (2013). Regulation of Adherens Junctions in Trabecular Meshwork Cells by Rac GTPase and their influence on Intraocular Pressure. J Ocul Biol 1. 10.13188/2334-2838.1000002.

Quinlan, A.R., and Hall, I.M. (2010). BEDTools: a flexible suite of utilities for comparing genomic features. Bioinformatics 26, 841–842. 10.1093/bioinformatics/btq033.

Raghunathan, V.K., Morgan, J.T., Park, S.A., Weber, D., Phinney, B.S., Murphy, C.J., and Russell, P. (2015). Dexamethasone Stiffens Trabecular Meshwork, Trabecular Meshwork Cells, and Matrix. Invest Ophthalmol Vis Sci 56, 4447–4459. 10.1167/iovs.15-16739.

Ramírez, F., Dündar, F., Diehl, S., Grüning, B.A., and Manke, T. (2014). deepTools: a flexible platform for exploring deep-sequencing data. Nucleic Acids Research 42, W187–W191. 10.1093/nar/gku365.

Rao, Z., Brunner, E., Giszas, B., Iyer-Bierhoff, A., Gerstmeier, J., Borner, F., Jordan, P.M., Pace, S., Meyer, K.P.L., Hofstetter, R.K., et al. (2023). Glucocorticoids regulate lipid mediator networks by reciprocal modulation of 15-lipoxygenase isoforms affecting inflammation resolution. Proc Natl Acad Sci U S A 120, e2302070120. 10.1073/pnas.2302070120.

Rauluseviciute, I., Riudavets-Puig, R., Blanc-Mathieu, R., Castro-Mondragon, J.A., Ferenc, K., Kumar, V., Lemma, R.B., Lucas, J., Cheneby, J., Baranasic, D., et al. (2024). JASPAR 2024: 20th anniversary of the open-access database of transcription factor binding profiles. Nucleic Acids Res 52, D174–D182. 10.1093/nar/gkad1059.

Robinson, M.D., McCarthy, D.J., and Smyth, G.K. (2010). edgeR: a Bioconductor package for differential expression analysis of digital gene expression data. Bioinformatics 26, 139–140. 10.1093/bioinformatics/btp616.

Rozsa, F.W., Reed, D.M., Scott, K.M., Pawar, H., Moroi, S.E., Kijek, T.G., Krafchak, C.M., Othman, M.I., Vollrath, D., Elner, V.M., and Richards, J.E. (2006). Gene expression profile of human trabecular meshwork cells in response to long-term dexamethasone exposure. Mol Vis 12, 125–141.

Saccuzzo, E.G., Youngblood, H.A., and Lieberman, R.L. (2023). Myocilin misfolding and glaucoma: A 20-year update. Prog Retin Eye Res 95, 101188. 10.1016/j.preteyeres.2023.101188.

Schipper, M., and Posthuma, D. (2022). Demystifying non-coding GWAS variants: an overview of computational tools and methods. Hum Mol Genet 31, R73–R83. 10.1093/hmg/ddac198.

Schoenfelder, S., and Fraser, P. (2019). Long-range enhancer-promoter contacts in gene expression control. Nat Rev Genet 20, 437–455. 10.1038/s41576-019-0128-0.

Schoenfelder, S., Javierre, B.M., Furlan-Magaril, M., Wingett, S.W., and Fraser, P. (2018). Promoter Capture Hi-C: High-resolution, Genome-wide Profiling of Promoter Interactions. J Vis Exp. 10.3791/57320.

Shah, R.N., Grzybowski, A.T., Cornett, E.M., Johnstone, A.L., Dickson, B.M., Boone, B.A., Cheek, M.A., Cowles, M.W., Maryanski, D., Meiners, M.J., et al. (2018). Examining the Roles of H3K4 Methylation States with Systematically Characterized Antibodies. Mol Cell 72, 162–177 e167. 10.1016/j.molcel.2018.08.015.

Shan, S., Wu, J., Cao, J., Feng, Y., Zhou, J., Luo, Z., Song, P., Rudan, I., and Global Health Epidemiology Research, G. (2024). Global incidence and risk factors for glaucoma: A systematic review and meta-analysis of prospective studies. J Glob Health 14, 04252. 10.7189/jogh.14.04252.

Shyam, R., Shen, X., Yue, B.Y., and Wentz-Hunter, K.K. (2010). Wnt gene expression in human trabecular meshwork cells. Mol Vis 16, 122–129.

Soneson, C., Love, M.I., and Robinson, M.D. (2015). Differential analyses for RNA-seq: transcript-level estimates improve gene-level inferences. F1000Res 4, 1521. 10.12688/f1000research.7563.2.

Soundararajan, A., Ghag, S.A., Vuda, S.S., Wang, T., and Pattabiraman, P.P. (2021). Cathepsin K Regulates Intraocular Pressure by Modulating Extracellular Matrix Remodeling and Actin-Bundling in the Trabecular Meshwork Outflow Pathway. Cells 10. 10.3390/cells10112864.

Soundararajan, A., Wang, T., Sundararajan, R., Wijeratne, A., Mosley, A., Harvey, F.C., Bhattacharya, S., and Pattabiraman, P.P. (2022). Multiomics analysis reveals the mechanical stress-dependent changes in trabecular meshwork cytoskeletal-extracellular matrix interactions. Front Cell Dev Biol 10, 874828. 10.3389/fcell.2022.874828.

Tamm, E.R. (2009). The trabecular meshwork outflow pathways: structural and functional aspects. Exp Eye Res 88, 648–655. 10.1016/j.exer.2009.02.007.

Tejwani, S., Machiraju, P., Nair, A.P., Ghosh, A., Das, R.K., Ghosh, A., and Sethu, S. (2020). Treatment of glaucoma by prostaglandin agonists and beta-blockers in combination directly reduces pro-fibrotic gene expression in trabecular meshwork. J Cell Mol Med 24, 5195–5204. 10.1111/jcmm.15172.

Tham, Y.C., Li, X., Wong, T.Y., Quigley, H.A., Aung, T., and Cheng, C.Y. (2014). Global prevalence of glaucoma and projections of glaucoma burden through 2040: a systematic review and meta-analysis. Ophthalmology 121, 2081–2090. 10.1016/j.ophtha.2014.05.013.

Uciechowski, P., and Dempke, W.C.M. (2020). Interleukin-6: A Masterplayer in the Cytokine Network. Oncology 98, 131–137. 10.1159/000505099.

van Koolwijk, L.M., Ramdas, W.D., Ikram, M.K., Jansonius, N.M., Pasutto, F., Hysi, P.G., Macgregor, S., Janssen, S.F., Hewitt, A.W., Viswanathan, A.C., et al. (2012). Common genetic determinants of intraocular pressure and primary open-angle glaucoma. PLoS genetics 8, e1002611. 10.1371/journal.pgen.1002611.

Vane, J.R., Bakhle, Y.S., and Botting, R.M. (1998). Cyclooxygenases 1 and 2. Annu Rev Pharmacol Toxicol 38, 97–120. 10.1146/annurev.pharmtox.38.1.97.

Vranka, J.A., Kelley, M.J., Acott, T.S., and Keller, K.E. (2015). Extracellular matrix in the trabecular meshwork: intraocular pressure regulation and dysregulation in glaucoma. Exp Eye Res 133, 112–125. 10.1016/j.exer.2014.07.014.

Wang, H., Duan, A., Zhang, J., Wang, Q., Xing, Y., Qin, Z., Liu, Z., and Yang, J. (2021). Glucocorticoid receptor wields chromatin interactions to tune transcription for cytoskeleton stabilization in podocytes. Commun Biol 4, 675. 10.1038/s42003-021-02209-8.

Wang, J., Qu, D., An, J., Yuan, G., and Liu, Y. (2017). Integrated microarray analysis provided novel insights to the pathogenesis of glaucoma. Mol Med Rep 16, 8735–8746. 10.3892/mmr.2017.7711.

Wang, Y., Li, X., and Hu, H. (2014). H3K4me2 reliably defines transcription factor binding regions in different cells. Genomics 103, 222–228. 10.1016/j.ygeno.2014.02.002.

Wang, Y., Song, C., Zhao, J., Zhang, Y., Zhao, X., Feng, C., Zhang, G., Zhu, J., Wang, F., Qian, F., et al. (2023). SEdb 2.0: a comprehensive super-enhancer database of human and mouse. Nucleic Acids Res 51, D280–D290. 10.1093/nar/gkac968.

Watanabe, M., Ida, Y., Ohguro, H., Ota, C., and Hikage, F. (2021). Establishment of appropriate glaucoma models using dexamethasone or TGFbeta2 treated three-dimension (3D) cultured human trabecular meshwork (HTM) cells. Sci Rep 11, 19369. 10.1038/s41598-021-98766-3.

Webber, H.C., Bermudez, J.Y., Millar, J.C., Mao, W., and Clark, A.F. (2018). The Role of Wnt/beta-Catenin Signaling and K-Cadherin in the Regulation of Intraocular Pressure. Invest Ophthalmol Vis Sci 59, 1454–1466. 10.1167/iovs.17-21964.

Weinreb, R.N., Leung, C.K., Crowston, J.G., Medeiros, F.A., Friedman, D.S., Wiggs, J.L., and Martin, K.R. (2016). Primary open-angle glaucoma. Nat Rev Dis Primers 2, 16067. 10.1038/nrdp.2016.67.

Whyte, W.A., Orlando, D.A., Hnisz, D., Abraham, B.J., Lin, C.Y., Kagey, M.H., Rahl, P.B., Lee, T.I., and Young, R.A. (2013). Master transcription factors and mediator establish super-enhancers at key cell identity genes. Cell 153, 307–319. 10.1016/j.cell.2013.03.035.

Wils, J., van Geuns, H., Stoot, J., Bergmans, M., Boschma, F., Bron, H., Degen, J., Erdkamp, F., van Erp, J., Haest, J., et al. (1999). Cyclophosphamide, epirubicin and cisplatin (CEP) versus epirubicin plus cisplatin (EP) in stage Ic-IV ovarian cancer: a randomized phase III trial of the Gynecologic Oncology Group of the Comprehensive Cancer Center Limburg. Anticancer Drugs 10, 257–261. 10.1097/00001813-199903000-00001.

Wingett, S., Ewels, P., Furlan-Magaril, M., Nagano, T., Schoenfelder, S., Fraser, P., and Andrews, S. (2015). HiCUP: pipeline for mapping and processing Hi-C data. F1000Res 4, 1310. 10.12688/f1000research.7334.1.

Winkler, N.S., and Fautsch, M.P. (2014). Effects of prostaglandin analogues on aqueous humor outflow pathways. J Ocul Pharmacol Ther 30, 102–109. 10.1089/jop.2013.0179.

Wu, J., Wang, C., Sun, S., Ren, T., Pan, L., Liu, H., Hou, S., Wu, S., Yan, X., Zhang, J., et al. (2024). Single-cell transcriptomic Atlas of aging macaque ocular outflow tissues. Protein Cell 15, 594–611. 10.1093/procel/pwad067.

Wu, Y., Yang, Y., Gu, H., Tao, B., Zhang, E., Wei, J., Wang, Z., Liu, A., Sun, R., Chen, M., et al. (2020). Multi-omics analysis reveals the functional transcription and potential translation of enhancers. Int J Cancer 147, 2210–2224. 10.1002/ijc.33132.

Xu, Z., Hysi, P., and Khawaja, A.P. (2021). Genetic Determinants of Intraocular Pressure. Annu Rev Vis Sci 7, 727–746. 10.1146/annurev-vision-031021-095225.

Yemanyi, F., Vranka, J., and Raghunathan, V.K. (2020). Glucocorticoid-induced cell-derived matrix modulates transforming growth factor beta2 signaling in human trabecular meshwork cells. Sci Rep 10, 15641. 10.1038/s41598-020-72779-w.

Yu, G., Wang, L.G., Han, Y., and He, Q.Y. (2012). clusterProfiler: an R package for comparing biological themes among gene clusters. OMICS 16, 284–287. 10.1089/omi.2011.0118.

Zhao, Y., Zhang, F., Pan, Z., Luo, H., Liu, K., and Duan, X. (2019). LncRNA NR_003923 promotes cell proliferation, migration, fibrosis, and autophagy via the miR-760/miR-215-3p/IL22RA1 axis in human Tenon’s capsule fibroblasts. Cell Death Dis 10, 594. 10.1038/s41419-019-1829-1.

Zhu, Y., Sun, L., Chen, Z., Whitaker, J.W., Wang, T., and Wang, W. (2013). Predicting enhancer transcription and activity from chromatin modifications. Nucleic Acids Res 41, 10032–10043. 10.1093/nar/gkt826.

