## Supplemental Data for "Chromatin dynamics identifies 78 genes at loci associated with elevated intraocular pressure and primary open-angle glaucoma"

**Supplemental Figure legends**

**Fig. S1. [A]** Expression of select single cell markers from (PMID: 32341164) in bulk total RNAseq from three TM-derived cell cultures. **[B]** Mean expression per 50kb windows separated by epigenetic mark peak density (Low = bottom quartile, mid = middle two quartiles, high = top quartile). Note that two marks, CTCF and H3K4me3, have distributions which make it impossible to identify three distinct groups in these cases, low represents the bottom three quartiles and high represents the top quartile. Error bars represent standard deviation, ** $p < 0.01$; **** $p < 0.0001$; ns = not significant. **[C]** Gene expression based on distance from transcription start site to nearest super enhancer. Bar represents median, red dot represent mean. *** $p < 0.001$. **[D]** GO terms enriched among genes overlapping TM super enhancers. Tree branches group terms based on semantic similarity. **[E]** Number of samples from SEdb 2.0 (PMID: 36318264) with a super enhancer overlapping a TM super enhancer. **[F]** Reactome pathways enriched among genes with promoter loops conserved across at least 10 tissues and TM cells. **[G]** GO terms enriched among expressed genes with CTCF-anchored loops in TM.

**Fig. S2.** Global transcriptional patterns by patient line and treatment status captured by principal components **[A]** PC1 vs PC2, and **[B]** PC2 vs PC3. **[C]** Normalized expression of genes that respond to glucocorticoid receptor (GR) signaling before and after dexamethasone treatment. **[D]** Overview of pipeline used to identify putative enhancer RNA (eRNA). **[E]** Expression changes for putative eRNA. Select eRNA are annotated with the identity of the closest gene transcription start site. Positive fold change values indicate eRNA with higher expression in dexamethasone treated samples; negative fold change values indicate eRNA with lower expression in dexamethasone treatment. **[F]** Stranded total RNA-seq data and enhancer locations at the *ALOX15B* locus including a putative eRNA upstream of the gene TSS.

**Fig. S3**. **[A]** $\text{Log}_2$ signal change detected in H3K4me2 in 50kb genomic windows that shift significantly towards B, significantly towards A, or that do not change. **[B]** Contact map of the *ZBTB16* locus with the *ZBTB16* gene highlighted in red. Lower and upper triangles indicate contacts in control and dexamethasone treated samples, respectively. Contact loops in red on the left edge are unique to the control condition; contact loops in blue on the upper edge are unique to the treated condition. Enrichment of **[C]** Reactome pathways and **[D]** GO biological process terms among genes with predicted binding of CEBPD in their promoter region in dexamethasone treated samples only. Enrichment of **[E]** Reactome pathways and **[F]** GO biological process terms among genes with predicted binding of Jun in their promoter region in control samples only.

**Fig S4. [A]** KEGG AGE-RAGE pathway, color scale represents $\log_2$ change in expression after dexamethasone treatment. **[B]** KEGG TNF Signaling pathway, color scale represents $\log_2$ change in expression after dexamethasone treatment. Epigenetic and chromatin changes as well as IOP/POAG GWAS variants near **[C]** FRMD8, **[D]** CREB5, **[E]** IL6, and **[F]** ALOX15B.

**Fig. S5. [A]** Count of tissues from (PMID: 31501517) overlapping each POAG lead SNP-Promoter pair observed in TM cells. **[B]** Count of IOP and POAG lead variants and variants in LD (shown in parenthesis) overlapping a SE in each dataset. **[C]** Percent of samples from SEdb 2.0 (n=1,739) overlapping each super enhancer containing a lead IOP SNP.

#### Supplemental Figure 1

A

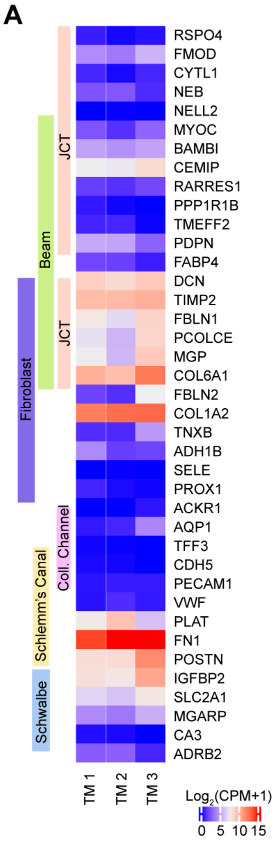


B

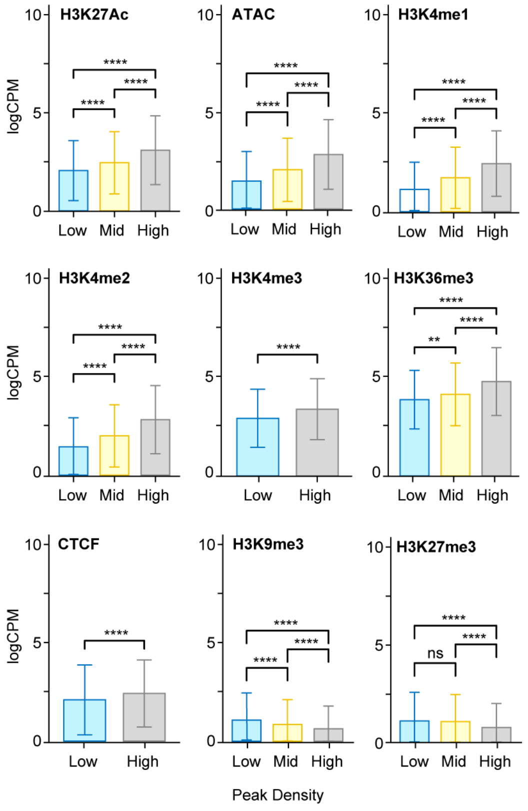


C

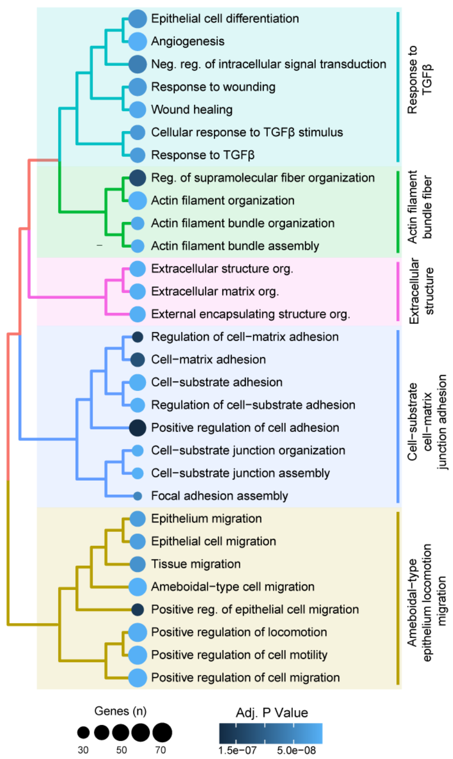


D

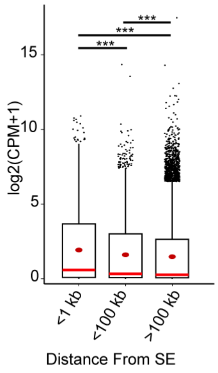


E

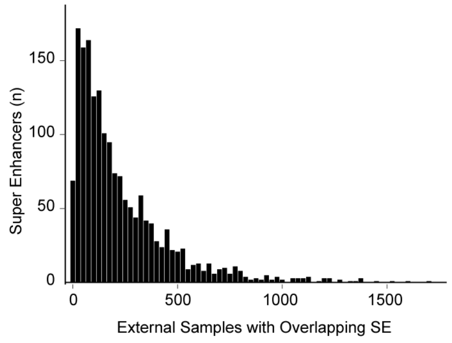


F

| GO Biological Process Term | Set Size | Hits | adj. *P* |
|---|---|---|---|
| Regulation of cellular component organization | 2433 | 94 | $7.97 \times 10^{-6}$ |
| Cellular response to stress | 1830 | 72 | $3.96 \times 10^{-4}$ |
| Negative regulation of cellular process | 5626 | 167 | $6.46 \times 10^{-4}$ |
| Chromatin organization | 884 | 43 | $7.63 \times 10^{-4}$ |
| Negative regulation of biological process | 5834 | 171 | $9.65 \times 10^{-4}$ |
| Positive regulation of biological process | 6264 | 180 | $1.53 \times 10^{-3}$ |
| Positive regulation of celullar process | 5920 | 172 | $1.60 \times 10^{-3}$ |
| Chromatin remodeling | 725 | 37 | $1.69 \times 10^{-3}$ |
| Organelle organization | 3594 | 116 | $1.89 \times 10^{-3}$ |
| Regulation of cell cycle | 1087 | 47 | $5.84 \times 10^{-3}$ |
| Cell cycle | 1663 | 63 | $9.06 \times 10^{-3}$ |
| DNA damage response | 908 | 41 | $9.33 \times 10^{-3}$ |
| Macromolecule modification | 3030 | 99 | $9.41 \times 10^{-3}$ |

G

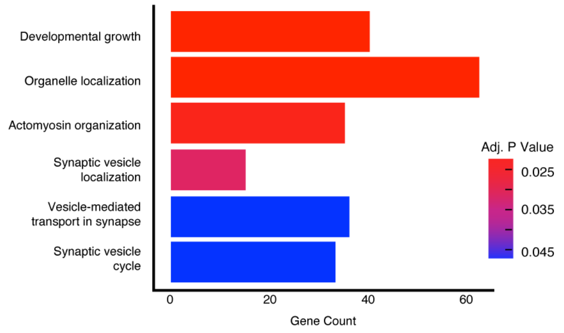

#### Supplemental Figure 2

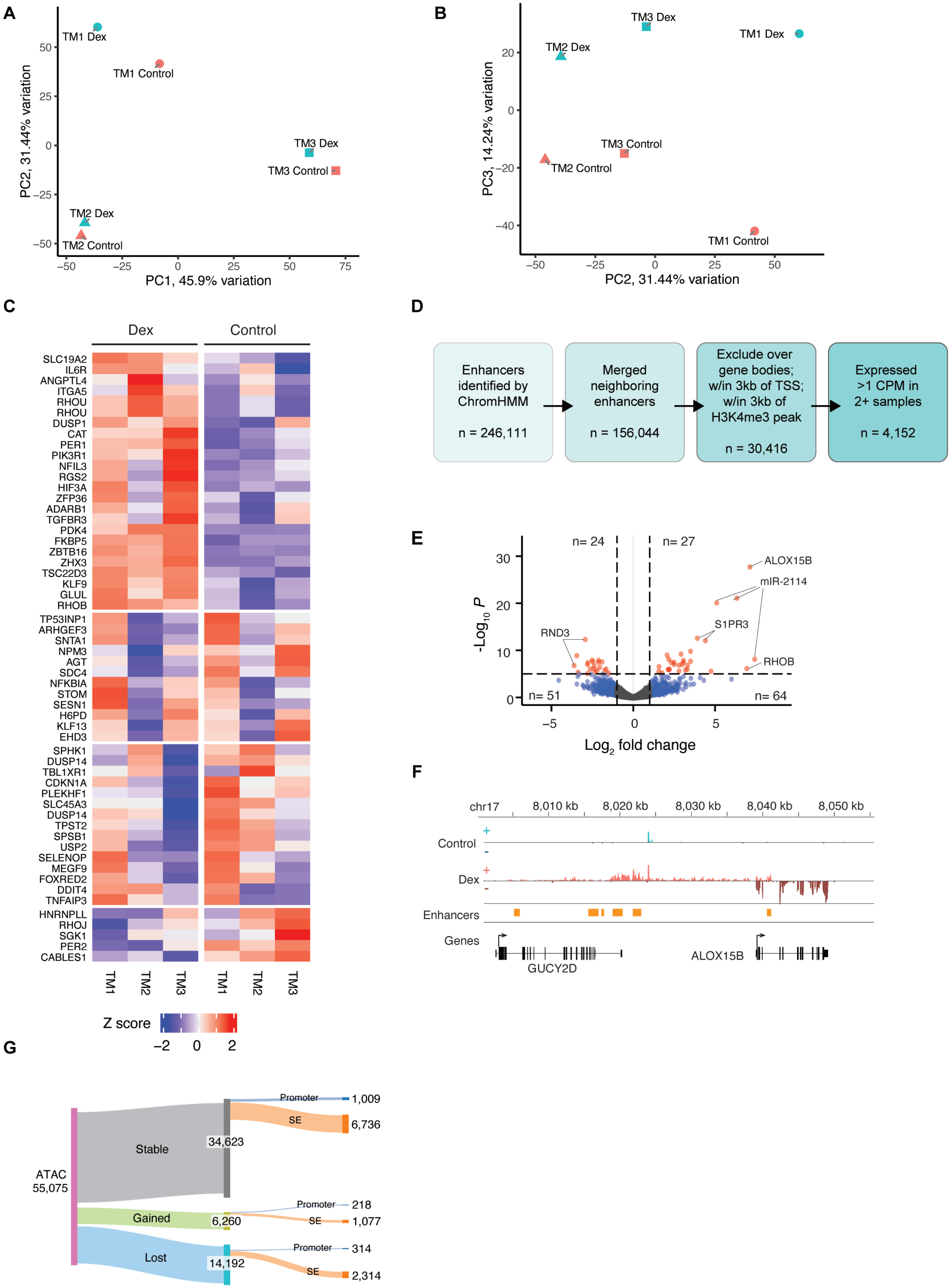

#### Supplemental Figure 3

A

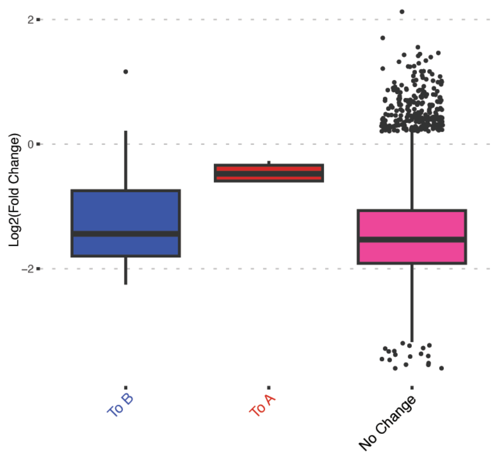


B

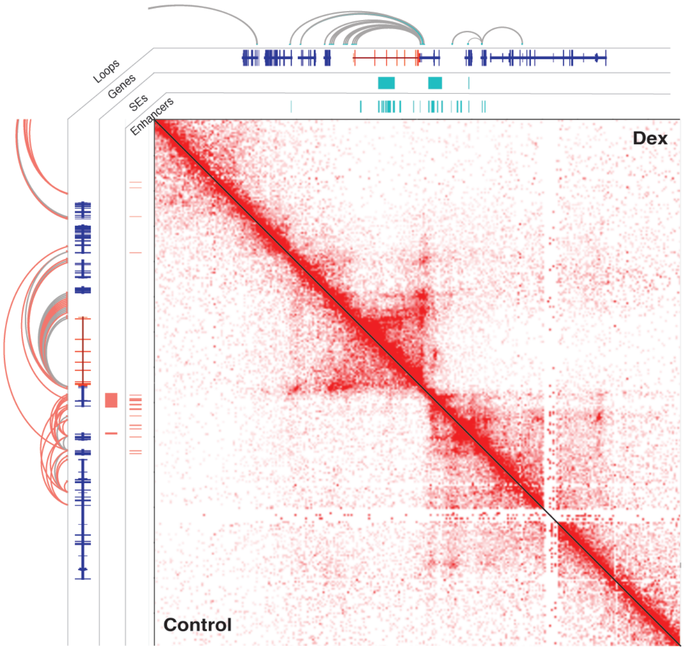


C

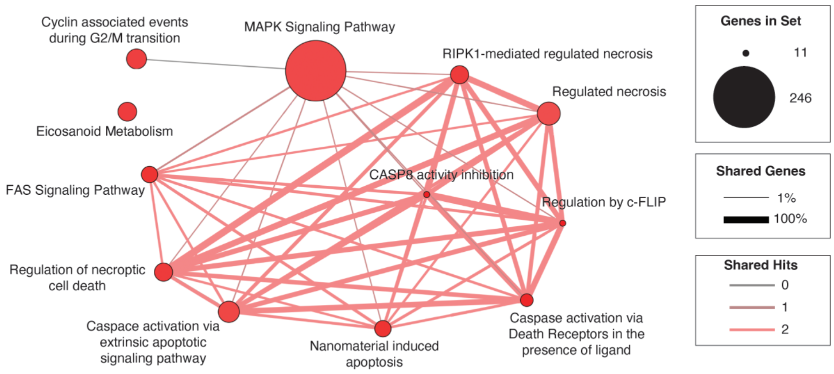


D

| GO BP Term | Term Size | Hit Size | *p* value |
|---|---|---|---|
| Organonitrogen compound metabolic process | 7,085 | 50 | $9.75 \times 10^{-5}$ |
| Mononuclear cell migration | 96 | 4 | 0.000975 |
| Glomerular mesangial cell proliferation | 11 | 2 | 0.0011 |
| Protein metabolic process | 5,979 | 41 | 0.00151 |
| Response to oxygen-containing compound | 1,592 | 16 | 0.00208 |
| Exocytosis | 901 | 11 | 0.00253 |
| Response to nitrogen compound | 1,067 | 12 | 0.00322 |
| Myeloid leukocyte migration | 220 | 5 | 0.00335 |
| Response to organic substance | 3,181 | 25 | 0.00345 |
| Negative regulation of biological process | 5,444 | 37 | 0.00364 |
| Leukocyte chemotaxis | 229 | 5 | 0.00397 |
| Extracellular structure organization | 454 | 7 | 0.00471 |
| Response to drug | 985 | 11 | 0.00506 |
| Symbiont process | 1,002 | 11 | 0.00578 |
| Macromolecule metabolic process | 9,610 | 56 | 0.00618 |

E

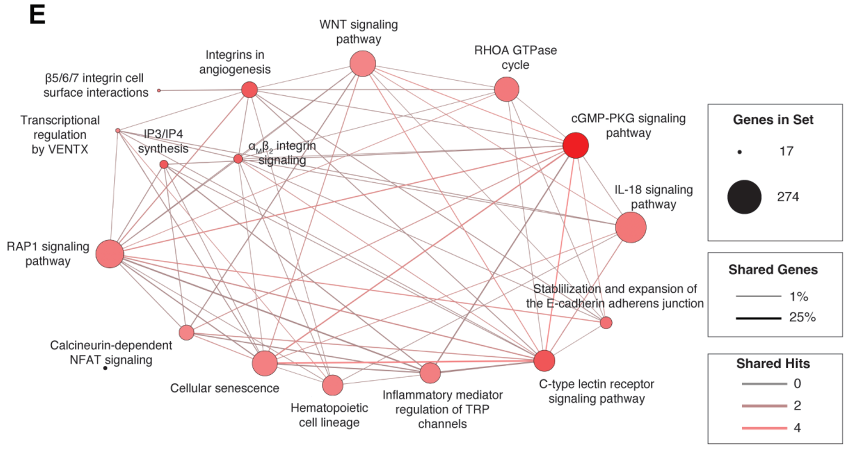


F

| GO BP Term | Term Size | Hit Size | *p* value |
|---|---|---|---|
| Chemokine production | 90 | 6 | $6.33 \times 10^{-5}$ |
| Cell surface receptor signaling pathway | 3,127 | 42 | $9.82 \times 10^{-5}$ |
| Cell-cell signaling by WNT | 513 | 13 | 0.000148 |
| Interleukin-8 production | 83 | 5 | 0.000423 |
| Actin cytoskeleton organization | 691 | 14 | 0.000771 |
| Interleukin-10 production | 57 | 4 | 0.000918 |
| Regulation of signaling | 3,492 | 42 | 0.00108 |
| Interleukin-1 production | 109 | 5 | 0.00145 |
| Tube development | 996 | 17 | 0.00147 |
| Interleukin-18 production | 9 | 2 | 0.00199 |
| Cell junction organization | 315 | 8 | 0.00282 |
| Apoptotic signaling pathway | 576 | 11 | 0.00452 |
| Programmed cell death | 2,054 | 26 | 0.00612 |
| Epithelium migration | 297 | 7 | 0.00762 |
| Regulation of cell adhesion | 713 | 12 | 0.00926 |

### Supplemental Figure 4

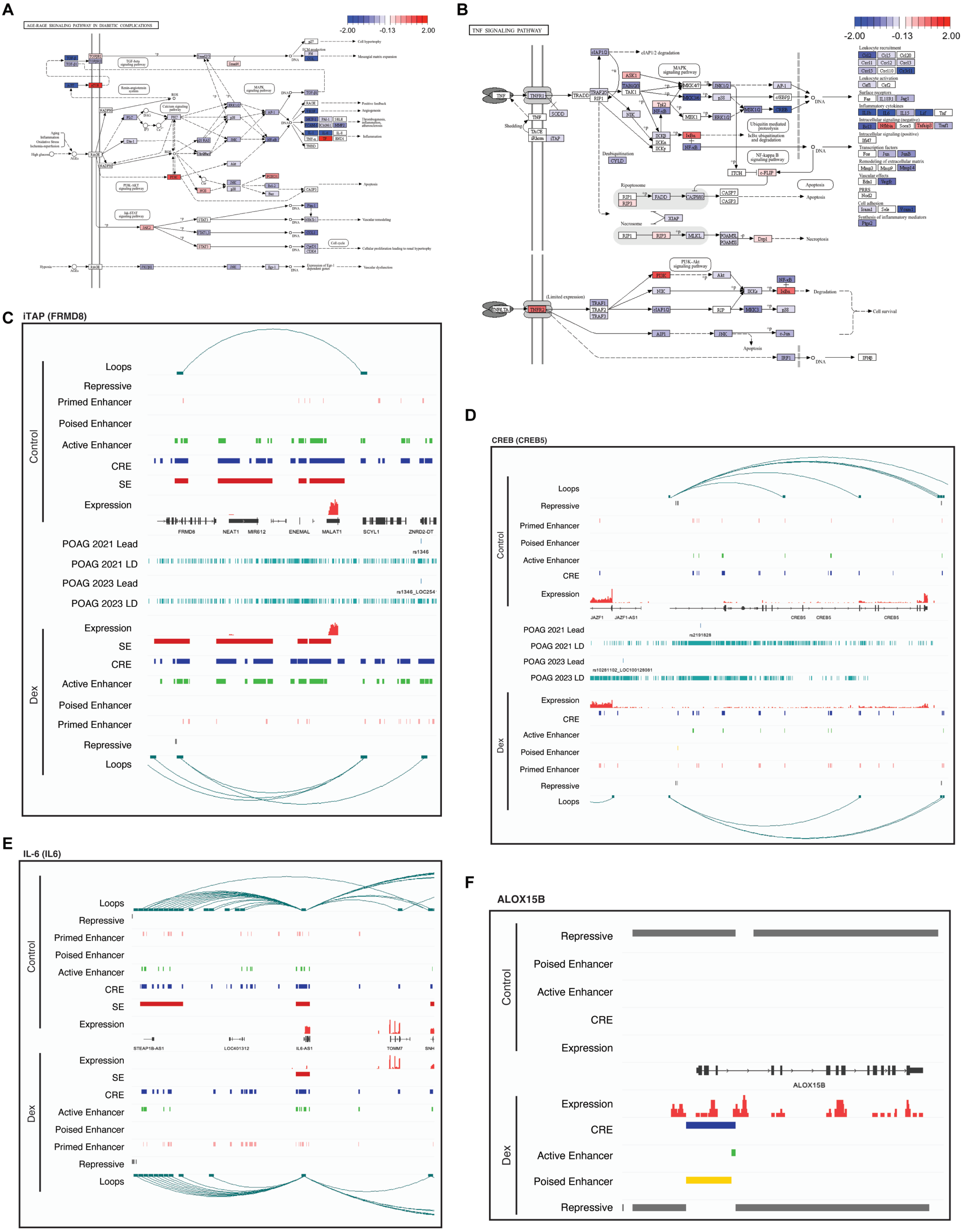

#### Supplemental Figure 5

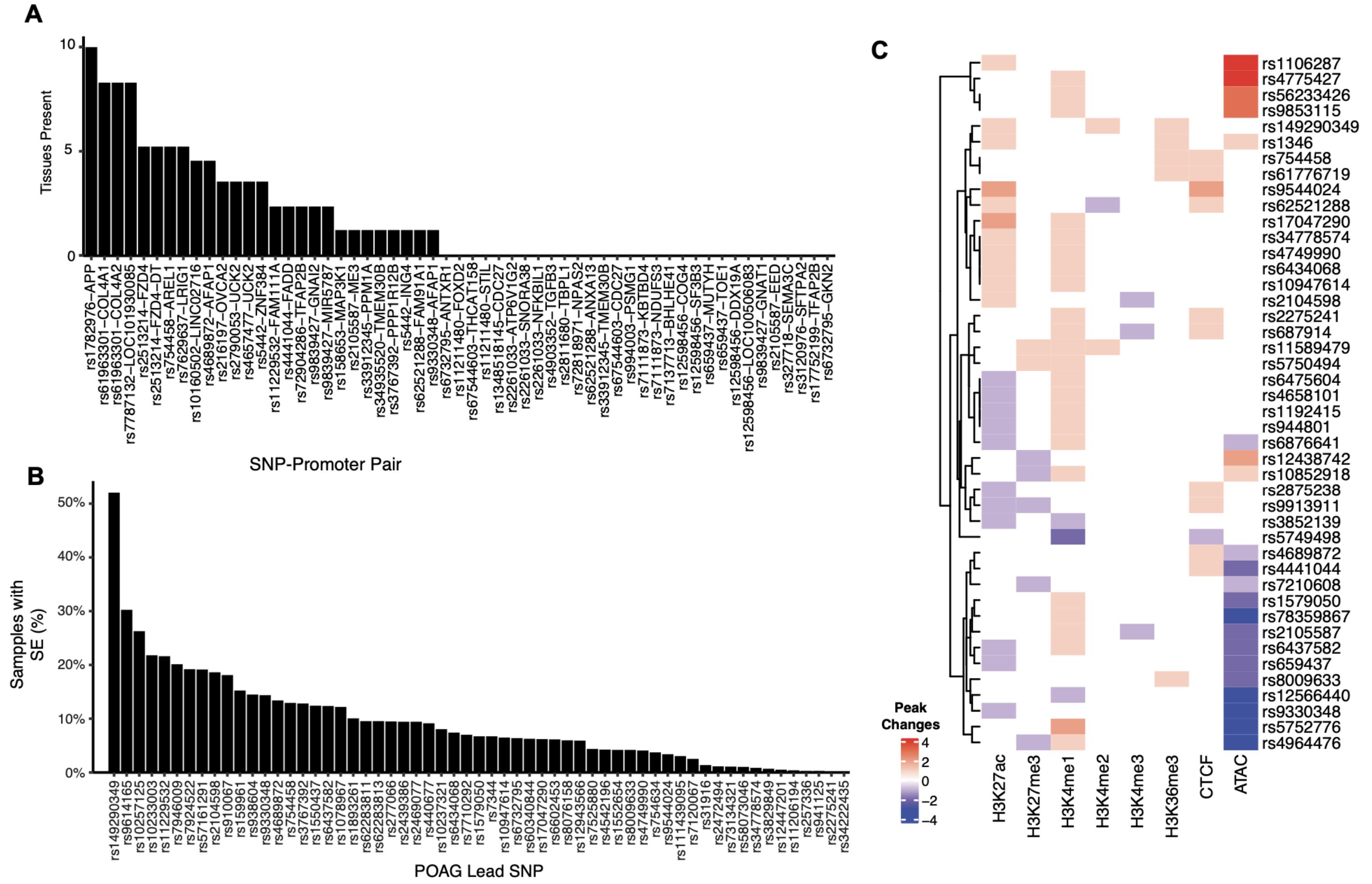

**Table S1. Target gene identified  Target gene identified for GWAS SNPs**

*POAG GWAS SNPs from Gharahkhani et al., 2021*

**Target gene for dexamethasone control**

| Lead variant | Loci | Chr | Chromosome position (hg38) | ID | Promoter target gene | Promoter length | Distance of variant from promoter |
|---|---|---|---|---|---|---|---|
| rs2790053 | TMCO1 | chr1 | 165768467 | ENSG00000283936 | MIR3658 | 359 | 139456 |
| rs2627761 | PNPT1/EFEMP1 | chr2 | 55705879 | ENSG00000207798 | MIR216A | 839 | 283907 |
| rs72904286 | TFAP2B/PKHD1 | chr6 | 51550124 | ENSG00000008196 | TFAP2B | 839 | -731215 |
| rs62521288 | FBXO32 | chr8 | 123542077 | ENSG00000104537 | ANXA13 | 719 | 195905 |
| rs62521288 | FBXO32 | chr8 | 123542077 | ENSG00000176853 | FAM91A1 | 719 | 226503 |
| rs33912345 | SIX6 | chr14 | 60509819 | ENSG00000182107 | TMEM30B | 598 | 772189 |
| rs754458 | LTBP2/AREL1 | chr14 | 74618126 | ENSG00000119682 | AREL1 | 826 | 94958 |
| rs4903352 | TTLL5 | chr14 | 75905315 | ENSG00000119699 | TGFB3 | 1500 | 78743 |
| rs13050568 | LOC101928435/PSMG1 | chr21 | 39034705 | ENSG00000157557 | ETS2 | 860 | -228669 |

**Targte gene dexamethasone treatment**

| Lead variant | Loci | Chr | Chromosome position (hg38) | ID | Promoter target gene | Promoter length | Distance of variant from promoter |
|---|---|---|---|---|---|---|---|
| rs2790053 | TMCO1 | chr1 | 165768467 | ENSG00000143179 | UCK2 | 859 | 59295 |
| rs62521288 | FBXO32 | chr8 | 123542077 | ENSG00000104537 | ANXA13 | 719 | 195905 |
| rs62521288 | FBXO32 | chr8 | 123542077 | ENSG00000176853 | FAM91A1 | 719 | 226503 |
| rs7111873 | RAPSN | chr11 | 47447887 | ENSG00000123444 | KBTBD4 | 479 | 131238 |
| rs7111873 | RAPSN | chr11 | 47447887 | ENSG00000025434 | NR1H3 | 1082 | -198567 |
| rs33912345 | SIX6 | chr14 | 60509819 | ENSG00000182107 | TMEM30B | 598 | 772189 |
| rs754458 | LTBP2/AREL1 | chr14 | 74618126 | ENSG00000119682 | AREL1 | 826 | 94958 |

**Target gene resource**

| Lead variant | Loci | Chr | Chromosome position (hg38) | ID | Promoter target gene | Promoter length | Distance of variant from promoter |
|---|---|---|---|---|---|---|---|
| rs2790053 | TMCO1 | chr1 | 165768467 | ENSG00000143179 | UCK2 | 859 | 59295 |
| rs2790053 | TMCO1 | chr1 | 165768467 | ENSG00000283936 | MIR3658 | 359 | 139456 |
| rs2627761 | PNPT1/EFEMP1 | chr2 | 55705879 | ENSG00000207548 | MIR217 | 359 | 277515 |
| rs6732795 | ANTXR1 | chr2 | 69184385 | ENSG00000183607 | GKN2 | 719 | -230920 |
| rs6732795 | ANTXR1 | chr2 | 69184385 | ENSG00000169604 | ANTXR1 | 599 | -171133 |
| rs72904286 | TFAP2B/PKHD1 | chr6 | 51550124 | ENSG00000008196 | TFAP2B | 839 | -731215 |
| rs62521288 | FBXO32 | chr8 | 123542077 | ENSG00000104537 | ANXA13 | 719 | 195905 |
| rs62521288 | FBXO32 | chr8 | 123542077 | ENSG00000176853 | FAM91A1 | 719 | 226503 |
| rs33912345 | SIX6 | chr14 | 60509819 | ENSG00000182107 | TMEM30B | 598 | 772189 |
| rs754458 | LTBP2/AREL1 | chr14 | 74618126 | ENSG00000119682 | AREL1 | 826 | 94958 |
| rs4903352 | TTLL5 | chr14 | 75905315 | ENSG00000119699 | TGFB3 | 1500 | 78743 |
| rs33912345 | SIX6 | chr14 | 60509819 | ENSG00000100614 | PPM1A | 359 | -263814 |
| rs216197 | SMG6 | chr17 | 2298650 | ENSG00000262664 | OVCA2 | 968 | -256467 |
| rs1348518145 | NPEPPS | chr17 | 47617876 | ENSG00000004897 | CDC27 | 479 | -428147 |

*POAG GWAS SNPs from Han et al., 2023*

| Lead variant | Loci | Chr | Chromosome position (hg38) | ID | Promoter target gene | Promoter length | Distance of variant from promoter |
|---|---|---|---|---|---|---|---|
| rs659437 | AKR1A1 | chr1 | 45571722 | ENSG00000132781 | MUTYH | 1079 | -231129 |
| rs7525880 | BCAR3 | chr1 | 93566431 | ENSG00000211575 | MIR760 | 119 | 280149 |
| rs4657477 | TMCO1 | chr1 | 165766938 | ENSG00000283936 | MIR3658 | 359 | 140985 |
| rs2627761 | PNPT1 | chr2 | 55705879 | ENSG00000207798 | MIR216A | 839 | 283907 |
| rs72818971 | NPAS2 | chr2 | 100986287 | ENSG00000170485 | NPAS2 | 119 | -166483 |
| rs9839427 | SEMA3B | chr3 | 50264152 | ENSG00000114349 | GNAT1 | 239 | -72471 |
| rs9839427 | SEMA3B | chr3 | 50264152 | ENSG00000114353 | GNAI2 | 1266 | -36867 |
| rs7629637 | KBTBD8 | chr3 | 66812641 | ENSG00000144749 | LRIG1 | 239 | -311218 |
| rs158653 | ANKRD55 | chr5 | 56282834 | ENSG00000095015 | MAP3K1 | 1002 | 532836 |
| rs2261033 | PRRC2A | chr6 | 31635814 | ENSG00000204463 | BAG6 | 1441 | 17487 |
| rs2261033 | PRRC2A | chr6 | 31635814 | ENSG00000265236 | SNORD84 | 599 | -94189 |
| rs2261033 | PRRC2A | chr6 | 31635814 | ENSG00000213760 | ATP6V1G2 | 479 | -89060 |

| | | | | | | | |
|---|---|---|---|---|---|---|---|
| rs2261033 | PRRC2A | chr6 | 31635814 | ENSG00000226979 | LTA | 599 | -63715 |
| rs2261033 | PRRC2A | chr6 | 31635814 | ENSG00000200816 | SNORA38 | 359 | -12656 |
| rs17752199 | PKHD1 | chr6 | 51542050 | ENSG00000008196 | TFAP2B | 839 | -723141 |
| rs7787132 | SEPTIN7 | chr7 | 35908275 | ENSG00000271122 | LOC101930085 | 843 | -212114 |
| rs327718 | SEMA3C | chr7 | 81210337 | ENSG00000075223 | SEMA3C | 479 | -287275 |
| rs3095122 | HSPA12A | chr10 | 116791252 | ENSG00000266782 | MIR3663 | 813 | 377177 |
| rs11229532 | LPXN | chr11 | 58574877 | ENSG00000166801 | FAM111A | 839 | 568395 |
| rs4441044 | ORAOV1 | chr11 | 69685595 | ENSG00000168040 | FADD | 935 | 517762 |
| rs2105587 | PRSS23 | chr11 | 86799140 | ENSG00000151376 | ME3 | 689 | -126113 |
| rs7941765 | ETS1 | chr11 | 128629105 | ENSG00000276176 | MIR6090 | 508 | -106269 |
| rs5442 | GNB3 | chr12 | 6845700 | ENSG00000111653 | ING4 | 719 | -181981 |
| rs7137713 | ITPR2 | chr12 | 26700989 | ENSG00000123095 | BHLHE41 | 239 | -575615 |
| rs61963301 | COL4A1 | chr13 | 110242338 | ENSG00000187498 | COL4A1 | 839 | 64788 |
| rs34935520 | SIX1 | chr14 | 60624683 | ENSG00000182107 | TMEM30B | 598 | 657325 |
| rs67544603 | ITGB3 | chr17 | 47250663 | ENSG00000263293 | THCAT158 | 839 | 73307 |
| rs1782976 | APP | chr21 | 25894555 | ENSG00000142192 | APP | 1098 | 277531 |
| rs9608740 | EMID1 | chr22 | 29224336 | ENSG00000100249 | C22orf31 | 719 | -161728 |

**Targte gene dexamethasone treatment**

| Lead variant | Loci | Chr | Chromosome position (hg38) | ID | Promoter target gene | Promoter length | Distance of variant from promoter |
|---|---|---|---|---|---|---|---|
| rs4657477 | TMCO1 | chr1 | 165766938 | ENSG00000143179 | UCK2 | 859 | 60824 |
| rs3767392 | PPP1R12B | chr1 | 202577973 | ENSG00000077157 | PPP1R12B | 359 | -229322 |
| rs9839427 | SEMA3B | chr3 | 50264152 | ENSG00000114353 | GNAI2 | 1266 | -36867 |
| rs7629637 | KBTBD8 | chr3 | 66812641 | ENSG00000144749 | LRIG1 | 239 | -311218 |
| rs2811680 | SLC2A12 | chr6 | 134056080 | ENSG00000028839 | TBPL1 | 710 | -103577 |
| rs3095122 | HSPA12A | chr10 | 116791252 | ENSG00000266782 | MIR3663 | 813 | 377177 |
| rs10160502 | SYT13 | chr11 | 45355265 | ENSG00000205106 | LINC02716 | 841 | 416363 |
| rs11229532 | LPXN | chr11 | 58574877 | ENSG00000166801 | FAM111A | 839 | 568395 |
| rs2105587 | PRSS23 | chr11 | 86799140 | ENSG00000151376 | ME3 | 689 | -126113 |
| rs2513214 | TMEM135 | chr11 | 87035111 | ENSG00000174804 | FZD4 | 938 | -79569 |
| rs61963301 | COL4A1 | chr13 | 110242338 | ENSG00000187498 | COL4A1 | 839 | 64788 |
| rs1782976 | APP | chr21 | 25894555 | ENSG00000142192 | APP | 1098 | 277531 |

**Target gene resource**

| Lead variant | Loci | Chr | Chromosome position (hg38) | ID | Promoter target gene | Promoter length | Distance of variant from promoter |
|---|---|---|---|---|---|---|---|
| rs659437 | AKR1A1 | chr1 | 45571722 | ENSG00000132781 | MUTYH | 1079 | -231129 |
| rs11211480 | TAL1 | chr1 | 47227548 | ENSG00000123473 | STIL | 239 | 86848 |
| rs11211480 | TAL1 | chr1 | 47227548 | ENSG00000186564 | FOXD2 | 359 | 208623 |
| rs4657477 | TMCO1 | chr1 | 165766938 | ENSG00000143179 | UCK2 | 859 | 60824 |
| rs4657477 | TMCO1 | chr1 | 165766938 | ENSG00000283936 | MIR3658 | 359 | 140985 |
| rs2627761 | PNPT1 | chr2 | 55705879 | ENSG00000207548 | MIR217 | 359 | 277515 |
| rs17047290 | EFEMP1 | chr2 | 55878799 | ENSG00000207798 | MIR216A | 839 | 110987 |
| rs6732795 | ANTXR1 | chr2 | 69184385 | ENSG00000183607 | GKN2 | 719 | -230920 |
| rs6732795 | ANTXR1 | chr2 | 69184385 | ENSG00000169604 | ANTXR1 | 599 | -171133 |
| rs9839427 | SEMA3B | chr3 | 50264152 | ENSG00000114353 | GNAI2 | 1266 | -36867 |
| rs7629637 | KBTBD8 | chr3 | 66812641 | ENSG00000144749 | LRIG1 | 239 | -311218 |
| rs4689872 | AFAP1 | chr4 | 7811565 | ENSG00000196526 | AFAP1 | 239 | 128597 |
| rs9330348 | AFAP1 | chr4 | 7882160 | ENSG00000196526 | AFAP1 | 239 | 58002 |
| rs158653 | ANKRD55 | chr5 | 56282834 | ENSG00000095015 | MAP3K1 | 1002 | 532836 |
| rs2261033 | PRRC2A | chr6 | 31635814 | ENSG00000204463 | BAG6 | 1441 | 17487 |
| rs2261033 | PRRC2A | chr6 | 31635814 | ENSG00000201785 | SNORD117 | 359 | -99053 |
| rs2261033 | PRRC2A | chr6 | 31635814 | ENSG00000265236 | SNORD84 | 599 | -94189 |
| rs2261033 | PRRC2A | chr6 | 31635814 | ENSG00000213760 | ATP6V1G2 | 479 | -89060 |
| rs2261033 | PRRC2A | chr6 | 31635814 | ENSG00000200816 | SNORA38 | 359 | -12656 |
| rs17752199 | PKHD1 | chr6 | 51542050 | ENSG00000008196 | TFAP2B | 839 | -723141 |
| rs2811680 | SLC2A12 | chr6 | 134056080 | ENSG00000028839 | TBPL1 | 710 | -103577 |
| rs7787132 | SEPTIN7 | chr7 | 35908275 | ENSG00000271122 | LOC101930085 | 843 | -212114 |
| rs3120976 | MAT1A | chr10 | 80265926 | ENSG00000185303 | SFTPA2 | 839 | -704973 |

| | | | | | | | |
|---|---|---|---|---|---|---|---|
| rs3095122 | HSPA12A | chr10 | 116791252 | ENSG00000266782 | MIR3663 | 813 | 377177 |
| rs11229532 | LPXN | chr11 | 58574877 | ENSG00000166801 | FAM111A | 839 | 568395 |
| rs2105587 | PRSS23 | chr11 | 86799140 | ENSG00000074266 | EED | 599 | -554351 |
| rs2105587 | PRSS23 | chr11 | 86799140 | ENSG00000151376 | ME3 | 689 | -126113 |
| rs2513214 | TMEM135 | chr11 | 87035111 | ENSG00000174804 | FZD4 | 938 | -79569 |
| rs5442 | GNB3 | chr12 | 6845700 | ENSG00000126746 | ZNF384 | 1304 | -155199 |
| rs7137713 | ITPR2 | chr12 | 26700989 | ENSG00000123095 | BHLHE41 | 239 | -575615 |
| rs61963301 | COL4A1 | chr13 | 110242338 | ENSG00000187498 | COL4A1 | 839 | 64788 |
| rs34935520 | SIX1 | chr14 | 60624683 | ENSG00000182107 | TMEM30B | 598 | 657325 |
| rs12598456 | IL34 | chr16 | 70642575 | ENSG00000261777 | LOC100506083 | 239 | -295697 |
| rs12598456 | IL34 | chr16 | 70642575 | ENSG00000103051 | COG4 | 479 | -118807 |
| rs67544603 | ITGB3 | chr17 | 47250663 | ENSG00000263293 | THCAT158 | 839 | 73307 |
| rs67544603 | ITGB3 | chr17 | 47250663 | ENSG00000004897 | CDC27 | 479 | -60934 |
| rs1782976 | APP | chr21 | 25894555 | ENSG00000142192 | APP | 1098 | 277531 |
| rs994003 | ERG | chr21 | 38665626 | ENSG00000183527 | PSMG1 | 580 | 518517 |

*IOP GWAS SNPs from Khawaja et al., 2018*

| Lead variant | Loci | Chr | Chromosome position (hg38) | ID | Promoter target gene | Promoter length | Distance of variant from promoter |
|---|---|---|---|---|---|---|---|
| rs116089225 | TMCO1 | chr1 | 165715441 | ENSG00000143179 | UCK2 | 859 | 112321 |
| rs116089225 | TMCO1 | chr1 | 165715441 | ENSG00000283936 | MIR3658 | 359 | 192482 |
| rs10918274 | TMCO1 | chr1 | 165745179 | ENSG00000283936 | MIR3658 | 359 | 162744 |
| rs6781336 | KBTBD8/LRIGI | chr3 | 66807626 | ENSG00000144749 | LRIG1 | 239 | -306203 |
| rs10036789 | PTCD2 | chr5 | 72400091 | ENSG00000083312 | TNPO1 | 1073 | 416676 |
| rs113985657 | EXOC2 | chr6 | 597203 | ENSG00000137273 | FOXF2 | 359 | 792475 |
| rs17752199 | PKHD1 | chr6 | 51542050 | ENSG00000008196 | TFAP2B | 839 | -723141 |
| rs327716 | HGF | chr7 | 81209661 | ENSG00000075223 | SEMA3C | 479 | -286599 |
| rs12356830 | NA | chr10 | 129032674 | ENSG00000108001 | EBF3 | 359 | 932324 |
| rs10838455 | SYT13 | chr11 | 45358223 | ENSG00000205106 | LINC02716 | 841 | 413405 |
| rs12923138 | ELMO3 | chr16 | 67199363 | ENSG00000172828 | CES3 | 360 | -238076 |
| rs17534001 | GNB1L/TXNRD2 | chr22 | 19854787 | ENSG00000184058 | TBX1 | 359 | -97867 |
| rs9608740 | EMID1 | chr22 | 29224336 | ENSG00000100249 | C22orf31 | 719 | -161728 |

**Targte gene dexamethasone treatment**

| Lead variant | Loci | Chr | Chromosome position (hg38) | ID | Promoter target gene | Promoter length | Distance of variant from promoter |
|---|---|---|---|---|---|---|---|
| rs6781336 | KBTBD8/LRIGI | chr3 | 66807626 | ENSG00000144749 | LRIG1 | 239 | -306203 |
| rs10036789 | PTCD2 | chr5 | 72400091 | ENSG00000083312 | TNPO1 | 1073 | 416676 |
| rs10838455 | SYT13 | chr11 | 45358223 | ENSG00000205106 | LINC02716 | 841 | 413405 |
| rs12923138 | ELMO3 | chr16 | 67199363 | ENSG00000172831 | CES2 | 599 | -264569 |

**Target gene resource**

| Lead variant | Loci | Chr | Chromosome position (hg38) | ID | Promoter target gene | Promoter length | Distance of variant from promoter |
|---|---|---|---|---|---|---|---|
| rs116089225 | TMCO1 | chr1 | 165715441 | ENSG00000283936 | MIR3658 | 359 | 192482 |
| rs6781336 | KBTBD8/LRIGI | chr3 | 66807626 | ENSG00000144749 | LRIG1 | 239 | -306203 |
| rs10036789 | PTCD2 | chr5 | 72400091 | ENSG00000083312 | TNPO1 | 1073 | 416676 |
| rs17752199 | PKHD1 | chr6 | 51542050 | ENSG00000008196 | TFAP2B | 839 | -723141 |
| rs10838455 | SYT13 | chr11 | 45358223 | ENSG00000205106 | LINC02716 | 841 | 413405 |
| rs7123436 | PTPRJ | chr11 | 47991932 | ENSG00000263693 | MIR3161 | 119 | 104832 |
| rs12923138 | ELMO3 | chr16 | 67199363 | ENSG00000159714 | ZDHHC1 | 359 | 217591 |
| rs12923138 | ELMO3 | chr16 | 67199363 | ENSG00000159713 | TPPP3 | 359 | 194284 |
| rs12923138 | ELMO3 | chr16 | 67199363 | ENSG00000172828 | CES3 | 360 | -238076 |
| rs35381200 | IL34 | chr16 | 70643165 | ENSG00000261777 | LOC100506083 | 239 | -296287 |
| rs35381200 | IL34 | chr16 | 70643165 | ENSG00000103051 | COG4 | 479 | -119397 |

*IOP GWAS SNPs from MacGregor et al., 2018*

| Lead variant | Loci | Chr | Chromosome position (hg38) | ID | Promoter target gene | Promoter length | Distance of variant from promoter |
|---|---|---|---|---|---|---|---|
| rs10918274 | TMCO1 | chr1 | 165745179 | ENSG00000283936 | MIR3658 | 359 | 162744 |

| | | | | | | | |
|---|---|---|---|---|---|---|---|
| rs11123857 | NPAS2 | chr2 | 100987350 | ENSG00000170485 | NPAS2 | 119 | -167546 |
| rs6781336 | LRIG1 | chr3 | 66807626 | ENSG00000144749 | LRIG1 | 239 | -306203 |
| rs10036789 | PTCD2 | chr5 | 72400091 | ENSG00000083312 | TNPO1 | 1073 | 416676 |
| rs113985657 | EXOC2 | chr6 | 597203 | ENSG00000137273 | FOXF2 | 359 | 792475 |
| rs17752199 | PKHD1 | chr6 | 51542050 | ENSG00000008196 | TFAP2B | 839 | -723141 |
| rs327716 | SEMA3C | chr7 | 81209661 | ENSG00000075223 | SEMA3C | 479 | -286599 |
| rs9608740 | EMID1 | chr22 | 29224336 | ENSG00000100249 | C22orf31 | 719 | -161728 |

**Targte gene dexamethasone treatment**

| Lead variant | Loci | Chr | Chromosome position (hg38) | ID | Promoter target gene | Promoter length | Distance of variant from promoter |
|---|---|---|---|---|---|---|---|
| rs6781336 | LRIG1 | chr3 | 66807626 | ENSG00000144749 | LRIG1 | 239 | -306203 |
| rs10036789 | PTCD2 | chr5 | 72400091 | ENSG00000083312 | TNPO1 | 1073 | 416676 |

**Target gene resource**

| Lead variant | Loci | Chr | Chromosome position (hg38) | ID | Promoter target gene | Promoter length | Distance of variant from promoter |
|---|---|---|---|---|---|---|---|
| rs6732795 | ANTXR1 | chr2 | 69184385 | ENSG00000183607 | GKN2 | 719 | -230920 |
| rs6732795 | ANTXR1 | chr2 | 69184385 | ENSG00000169604 | ANTXR1 | 599 | -171133 |
| rs6781336 | LRIG1 | chr3 | 66807626 | ENSG00000144749 | LRIG1 | 239 | -306203 |
| rs10036789 | PTCD2 | chr5 | 72400091 | ENSG00000083312 | TNPO1 | 1073 | 416676 |
| rs17752199 | PKHD1 | chr6 | 51542050 | ENSG00000008196 | TFAP2B | 839 | -723141 |
| rs2697920 | MYBPC3 | chr11 | 47349056 | ENSG00000275208 | MIR6745 | 355 | -168971 |
| rs7123436 | PTPRJ | chr11 | 47991932 | ENSG00000263693 | MIR3161 | 119 | 104832 |

*IOP GWAS SNPs from Gharahkhani et al., 2018*

| Lead variant | Loci | Chr | Chromosome position (hg38) | ID | Promoter target gene | Promoter length | Distance of variant from promoter |
|---|---|---|---|---|---|---|---|
| rs2790053 | LOC100147773 | chr1 | 165768467 | ENSG00000283936 | MIR3658 | 359 | 139456 |
| rs2627761 | PNPT1-EFEMP1 | chr2 | 55705879 | ENSG00000207798 | MIR216A | 839 | 283907 |
| rs72904286 | TFAP2B-PKHD1 | chr6 | 51550124 | ENSG00000008196 | TFAP2B | 839 | -731215 |
| rs62521288 | FBXO32 | chr8 | 123542077 | ENSG00000104537 | ANXA13 | 719 | 195905 |
| rs62521288 | FBXO32 | chr8 | 123542077 | ENSG00000176853 | FAM91A1 | 719 | 226503 |
| rs754458 | LTBP2-AREL1 | chr14 | 74618126 | ENSG00000119682 | AREL1 | 826 | 94958 |

**Targte gene dexamethasone treatment**

| Lead variant | Loci | Chr | Chromosome position (hg38) | ID | Promoter target gene | Promoter length | Distance of variant from promoter |
|---|---|---|---|---|---|---|---|
| rs2790053 | LOC100147773 | chr1 | 165768467 | ENSG00000143179 | UCK2 | 859 | 59295 |
| rs62521288 | FBXO32 | chr8 | 123542077 | ENSG00000104537 | ANXA13 | 719 | 195905 |
| rs62521288 | FBXO32 | chr8 | 123542077 | ENSG00000176853 | FAM91A1 | 719 | 226503 |
| rs7111873 | RAPSN | chr11 | 47447887 | ENSG00000123444 | KBTBD4 | 479 | 131238 |
| rs7111873 | RAPSN | chr11 | 47447887 | ENSG00000025434 | NR1H3 | 1082 | -198567 |

**Target gene resource**

| Lead variant | Loci | Chr | Chromosome position (hg38) | ID | Promoter target gene | Promoter length | Distance of variant from promoter |
|---|---|---|---|---|---|---|---|
| rs2790053 | LOC100147773 | chr1 | 165768467 | ENSG00000143179 | UCK2 | 859 | 59295 |
| rs2790053 | LOC100147773 | chr1 | 165768467 | ENSG00000283936 | MIR3658 | 359 | 139456 |
| rs2627761 | PNPT1-EFEMP1 | chr2 | 55705879 | ENSG00000207548 | MIR217 | 359 | 277515 |
| rs6732795 | ANTXR1 | chr2 | 69184385 | ENSG00000183607 | GKN2 | 719 | -230920 |
| rs6732795 | ANTXR1 | chr2 | 69184385 | ENSG00000169604 | ANTXR1 | 599 | -171133 |
| rs72904286 | TFAP2B-PKHD1 | chr6 | 51550124 | ENSG00000008196 | TFAP2B | 839 | -731215 |
| rs62521288 | FBXO32 | chr8 | 123542077 | ENSG00000104537 | ANXA13 | 719 | 195905 |
| rs62521288 | FBXO32 | chr8 | 123542077 | ENSG00000176853 | FAM91A1 | 719 | 226503 |
| rs754458 | LTBP2-AREL1 | chr14 | 74618126 | ENSG00000119682 | AREL1 | 826 | 94958 |
